# A 3D iPSC retina model reveals non-cell-autonomous and non-neuronal mechanism of photoreceptor degeneration in a lysosomal storage disorder

**DOI:** 10.1101/2025.07.10.664233

**Authors:** Jimin Han, Nathaniel Foley, Sonal Dalvi, Janet A. H. Tang, Amit Chatterjee, Lal Krishan Kumar, Chad A. Galloway, Kumar Singh, Yashoda Subedi, Kevin Ling, Alison Heffer, Richard T. Libby, Danielle S. W. Benoit, Anthony L. Cook, Vera Bonilha, Edward Schuchman, Jennifer J. Hunter, Ruchira Singh

## Abstract

Disruption of photoreceptor-retinal pigment epithelium (RPE) interface with loss of photoreceptor outer segments (POSs) in the retina is a pathological hallmark of several neurodegenerative and retinal diseases including lysosomal storage disorder’s like CLN3 disease. However, the retina is a functional composite *in vivo;* and *in vitro* stem cell models of retina that enable investigation of the photoreceptor-RPE interface in healthy and diseased retina are lacking. Here, we developed a 3D human pluripotent stem cell (hPSC)-derived retina model to investigate the photoreceptor-RPE interface in healthy and disease tissue. Using this 3D hPSC retina model, we demonstrated that the most common disease causing *CLN3* mutation (*CLN3*^Δ*ex7-8*^) leads to reduced levels of acid ceramidase (AC) and consequently altered sphingolipid metabolism and signaling and POS loss in CLN3 disease. Consistent with the 3D hPSC retina model, altered sphingolipid metabolism and signaling coincided with POS loss in a large animal model of CLN3 disease, CLN3 miniswine. Therapeutically, recombinant human acid ceramidase (rhAC) targeted both altered sphingolipid metabolism and retina degeneration in the CLN3 hPSC retina model and the CLN3 miniswine eye. These findings demonstrate a proof-of-concept that rhAC can rescue disease phenotype in a large animal model of CLN3 disease and suggest that rhAC could be a therapeutic approach for CLN3 disease.

**One Sentence Summary:** Acid ceramidase deficiency and consequently altered sphingolipid signaling promotes disease phenotype(s) in a lysosomal storage disorder, CLN3 disease.

## INTRODUCTION

Neuronal Ceroid lipofuscinoses (NCL) diseases, collectively referred to as Batten disease, encompass a group of hereditary neurodegenerative disorders that largely belong to the category of lysosomal storage disorders (LSD) (*1–4*). NCLs are the most common type of progressive juvenile neurodegeneration with a prevalence of 1:12,500 in some populations (*1–4*). NCLs share a variety of neurodegenerative symptoms reminiscent of dementia, as well as visual impairment, motor function decline, seizures, and shortened life expectancy (*1–4*).

CLN3 disease (also referred to as juvenile ceroid lipofuscinosis or JNCL) caused by mutations in *CLN3* gene is the most common form of NCLs (*5–8*). Vision loss in early childhood followed by legal blindness is seen in all patients with CLN3 disease (*9–13*). Neurological symptoms that follow vision loss in CLN3 disease include decline in speech and motor skills, cognitive and behavioral problems, seizures, and premature death by the third decade in life (44, 46–49). Although there are currently no treatments for CLN3 disease, the therapeutic pipeline for CLN3 disease includes mutation-specific approaches including Antisense Oligonucleotide (ASO) treatment that recently received FDA approval for an N=2 study, (*14, 15*) and AAV-based gene therapy approaches targeting the brain and the retina (NCT03770572, NCT05228145, NCT04737460). In addition, mutation-independent approaches such as targeting glycosphingolipid accumulation and neuroinflammation (Batten-/miglustat; NCT05174039) and activating peroxisome proliferator-activated receptor-alpha (PPAR-α) to target lipid metabolism (PLX-200/gemfibrozil; NCT04637282) are currently being pursued as treatment options for CLN3 disease. There are also several other therapeutic development efforts in progress for CLN3 disease that target specific cellular pathways including activation of TRPML1 channel and targeting c-Abl kinase – mediated activation of YAP1 pro-apoptotic signaling (*16, 17*).

Despite ongoing clinical trials, a major challenge for development of rational therapeutics for CLN3 disease is the lack of understanding of the underlying disease mechanism. *CLN3* is a ubiquitously expressed gene and CLN3 protein is predominantly localized to the lysosomal/endosomal membrane in the cell (*18, 19*). Consistently, several studies have indicated a role of CLN3 protein in regulation of endosomal-lysosomal homeostasis including lysosomal trafficking (*19–21*). In addition, CLN3 protein has also been linked to regulation of actin cytoskeleton, apoptosis, calcium signaling, lipid metabolism, mitochondrial homeostasis and osmoregulation (*7, 16, 17, 19, 20, 22–24*). Recent studies have also shown a cell autonomous function of CLN3 in non-neuronal cells, including in the brain (e.g., microglia and astrocytes) and the retina (retinal pigment epithelium or RPE) and suggested a role of these supporting cells in CLN3 disease neuropathology (*10, 25–27*). Overall, the aforementioned studies have significantly increased both our understanding of the diverse role of CLN3 gene/protein in cellular homeostasis and molecular and pathological changes in CLN3 disease.

However, the cellular and molecular driver(s) of CLN3 disease pathology in the retina and brain remains to be identified. This is partly due to lack of access to human retina and brain tissue for molecular studies in early-stage disease. Given that the earliest pathological manifestations of CLN3 disease affect the photoreceptor-retinal pigment epithelium (RPE) interface in the retina (*9, 28, 29*), a human cell model that enables investigation of the photoreceptor-RPE complex would be valuable for investigating CLN3 disease biology. Note that although human pluripotent stem cell (hPSCs) derived neural retina (retinal organoid/RO) and RPE monocultures have provided a powerful human platform for studying retinal diseases (*30–36*), the photoreceptor containing RO and RPE do not co-develop in the hPSC model of retinogenesis (*37, 38*).

To address the lack of hPSC-retina model, we integrated hPSC-derived RO and RPE into a hydrogel-based, engineered extracellular matrix (ECM) to generate a 3D hPSC RO-RPE model that allowed investigation of photoreceptor-RPE complex in CLN3 disease. By analyzing control versus *CLN3* mutant (*CLN3*^Δ*ex7-8*^) hPSC-RPE monoculture and hPSC-RO-RPE model, we identified *i)* an important role of acid ceramidase (AC)-mediated lysosomal sphingolipid metabolism in CLN3 disease pathobiology and *ii)* a cell autonomous role of the RPE in promoting POS loss/retina degeneration in CLN3 disease. Furthermore, we validated molecular and structural changes observed in the hPSC-RPE monoculture and 3D RO-RPE model of CLN3 disease using *i)* a large animal model of CLN3 disease, CLN3 miniswine with the orthologous human mutation (*CLN3*^Δ*ex7-8*^) *ii)* CLN3 disease patient donor eyes and *iii)* high-resolution retina imaging on the living eye of CLN3 disease patients. Ultimately, we tested the therapeutic impact of recombinant acid ceramidase in the 3D hPSC RO-RPE model of CLN3 disease and CLN3 miniswine eyes.

## RESULTS

The complete list of abbreviations used in this study are provided in ***Table S1***. Furthermore, unless otherwise specified, all references to the RO-RPE model refer to hPSC-derived model.

### Engineered extracellular matrix enables RO-RPE assembloid with photoreceptor-RPE interaction

In the *in vivo* retina, a specialized extracellular matrix, the interphotoreceptor matrix (IPM) supports the neural retina and RPE adhesion and promotes development and maturation of the photoreceptor-RPE complex (*39, 40*). To mimic the highly hydrophilic matrix metalloproteinase (MMP)-degradable hyaluronan-rich IPM (*41*) *in vitro*, we utilized poly(ethylene glycol) (PEG) hydrogels incorporating hyaluronan (100 µg/ml) (PEG-HA) crosslinked with a MMP-degradable peptide (*Figure 1a*). Furthermore, to emulate IPM occupying the space between neural retina and RPE *in vivo*, hPSC-derived neural retina organoid (RO) were encapsulated in PEG-HA and co-cultured with RPE monolayer on transwells (*Figure 1a*).

**Figure 1.**
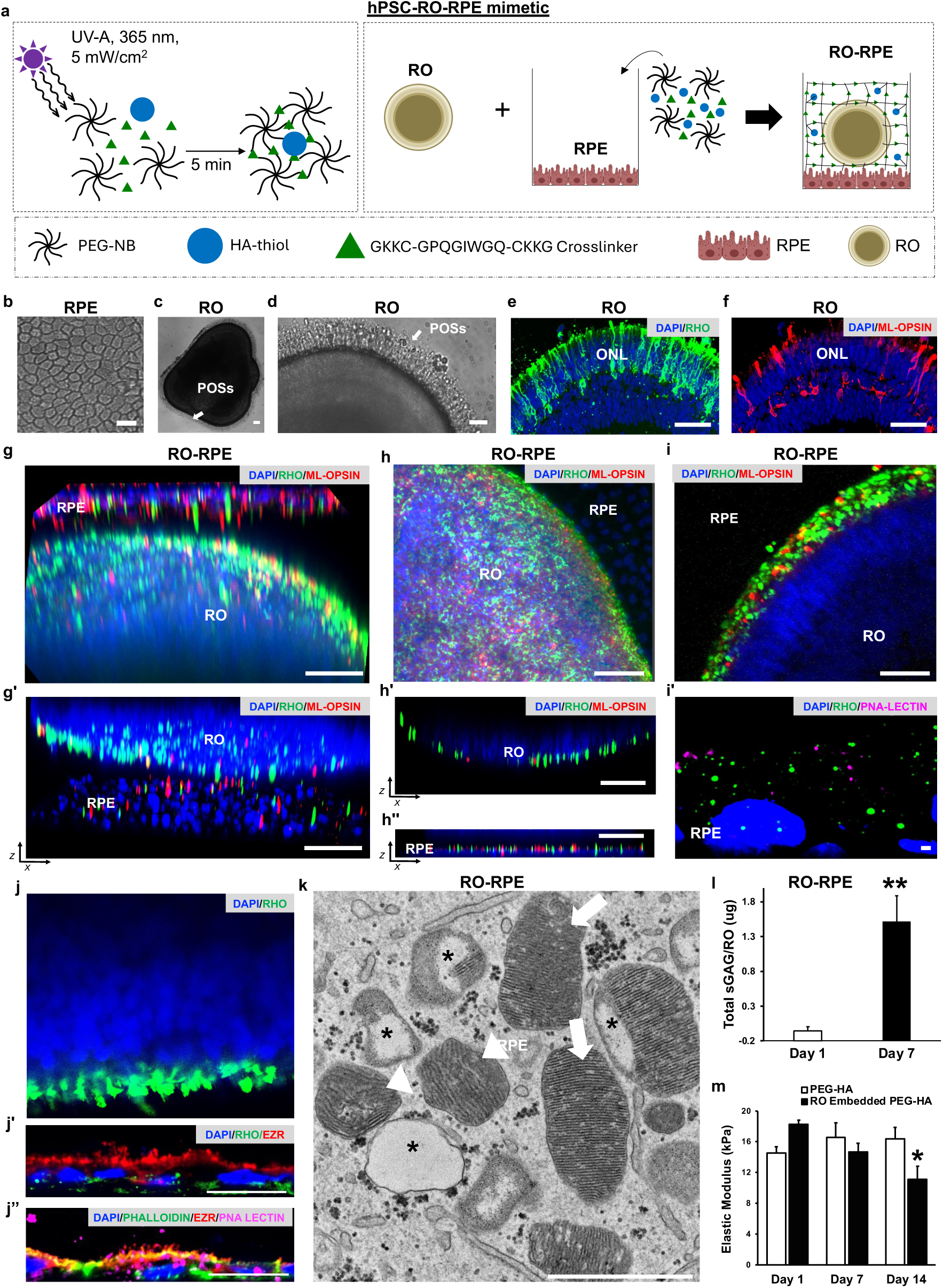
RO-RPE assembloid co-culture in engineered extracellular matrix promotes photoreceptor-RPE interaction. **a**) Schematic representation of the 3D hPSC RO-RPE assembloid development. RO was encapsulated in a solution containing PEG, hyaluronan, and MMP-degradable crosslinker over a monolayer of hPSC-RPE in transwells. **b-d)** Light microscopy images of RPE monolayer (b) and RO (c, d) monocultures prior to RO-RPE co-culture illustrating expected hexagonal morphology (b) and showing presence of hair-like projections (c, d) that are representative of photoreceptor outer segments (POSs) in RO (arrows). White arrow (c, d) denotes presence of POS in brightfield images. Scale bar = 50 µm. **e, f)** Immunofluorescence images of the stage 3 RO prior to use in RO-RPE co-culture illustrating the outer nuclear layer (ONL), the RHO^+^ rod outer segments (ROS) (green, e) and the ML-OPSIN^+^ cone outer segments (COS) (red, f). DAPI (blue; e, f) shows cell nuclei. Scale bar = 50 µm. **g**) Immunofluorescence images showing RHO^+^ ROS (green), ML-OPSIN^+^ COS (red), and DAPI^+^ cell nuclei (blue) in the RO and RPE monolayer of day 7 RO-RPE assembloid culture in 3D volume (g) and 3D orthogonal xz view (g’). Scale bar = 50 µm. **h, i)** Immunofluorescence images showing RHO^+^ ROS (green) and ML-OPSIN^+^ and PNA-LECTIN^+^ COS (red, magenta) in the RO layer of the day 7 RO-RPE assembloid culture in planar view (h, i) and orthogonal (xz) view at two different planes corresponding to RO (h’) and RPE monolayer (h’’). Scale bar = 50 µm. The immunofluorescence image of the cryosection of the RPE monolayer (i’) illustrates shed POSs (RHO^+^ ROS and PNA-LECTIN^+^ COS) above the RPE cells (arrow) and ingested POSs within the RPE cells (arrowhead). Cell nuclei (DAPI; blue) are shown in blue. Scale bar = 10 µm. **j**) Immunofluorescence images showing the spatial localization of RHO^+^ ROS in the RO layer (j) and RHO^+^ ROS, PNA-LECTIN^+^ COS and EZR-labeled apical microvilli and phalloidin stained actin cytoskeleton in the RPE monolayer of day 7 RO-RPE assembloid cultures (j, j’). Scale bar = 10 µm. **k**) Transmission electron microscopy image showing ingested POSs (ROS and COS) in an RPE cell of the day 7 RO-RPE assembloid culture. ROS (arrows) and COS (arrowhead) ingested by the RPE monolayer have the expected ultrastructure with parallel stacks of lobulated disks. The ROS and COS phagosomes are also seen at different stages of lysosomal degradation. Scale bar = 1 µm. **l**) Quantitative analysis of sulfated glycosaminoglycan (sGAG) deposition (l) of the RO tissue in the hydrogels. Increased deposition of sGAG is seen in the RO layer between day 1 and day 7 of the RO-RPE assembloid culture (l). **m**) Young’s modulus of the RO embedded PEG-HA is within the range of the elastic modulus reported for *in vivo* retina tissue (∼10-20 kPA) (*50*). The measurement of elastic modulus provided evidence of tissue remodeling with the RO embedded PEG-HA but not PEG-HA gels alone showing decrease in elastic modulus measurement with time in culture (day 1 to day 14). * p<0.05, ** p< 0.01. Biological replicate n ≧ 3 for all Figure 1 experiments.

A mature monolayer of RPE and stage 3 RO with presence of photoreceptor outer segments (POSs) were used in the RO-RPE assembloid culture (*Figure 1b-1f* and *S1a-S1l*). The mature RPE monolayer showed the characteristic hexagonal morphology (*Figure 1b*) and expected localization of apical microvilli protein, Ezrin (EZR), tight junction marker Zonula occludens-1 (ZO-1), and phalloidin-labeled F-actin cytoskeleton (*Figure S1a-S1c*). Similarly, consistent with published RO characterization at different stages of maturation (stage 1-3) (*42, 43*), the stage 3 ROs showed outer retina organization with presence of hair-like surface projections (POSs) (*Figure 1c, 1d*), a defined outer nuclear layer (ONL) with RCVRN-positive (RCVRN^+^) POSs and rhodopsin/RHO^+^ rod outer segments (ROSs) and M/L-opsin/ML-OPSIN^+^ and PNA Lectin^+^ cone outer segments (COSs) (*Figure 1e, 1f* and *S1d-S1l*). Furthermore consistent with prior RO characterization (*42, 43*), the stage 3 ROs showed presence of visual system homeobox 2-positive (VSX2)^+^ bipolar cells in the inner nuclear layer and cellular retinaldehyde-binding protein-positive (CRALBP^+^) Müller glia that extended both outwardly to form a ZO-1/EZR/Phalloidin^+^ external limiting membrane and inwardly into a poorly nucleated RO core with remnant tubulin beta 3 class III (TUBB3^+^) retinal ganglion cells and displaced RCVRN^+^, ML-OPSIN^+^ and RHO^+^ photoreceptor cells (*Figure S1d-S1l*).

Overall, immunocytochemical characterization of the RPE monolayer used for RO-RPE culture was consistent with a polarized epithelial barrier (*Figure 1b and S1a-S1c*). Furthermore, the stage 3 RO used in the RO-RPE culture showed degeneration and disorganization of the inner retina but displayed proper outer retina organization with presence of ONL, external limiting membrane and photoreceptor inner and outer segments (*Figure 1b-1f and S1d-S1l*). Therefore, we primarily focused on the characterization of the photoreceptor and RPE layer in the RO-RPE model.

Next, to critically evaluate the role of PEG-HA matrix and RPE monolayer on photoreceptor/RO properties, we utilized two distinct approaches. In the first approach, ONL/POS organization was compared in direct co-cultures of RO-RPE and RO monocultures in the absence of PEG-HA matrix (*Figure S2*). In the second approach, ONL/POS organization was compared between parallel cultures of PEG-HA RO-RPE and PEG-HA RO (*Figure 1 and S3*). For the latter, RO encapsulated in PEG-HA were cultured as monocultures in the absence of RPE. The RO monoculture and RO-RPE co-cultures in both conditions (direct co-culture and PEG-HA model) were maintained for 7 days.

Immunocytochemical characterization of RO cryosections in parallel cultures of RO and RO-RPE in the direct co-culture model did not show a positive impact of RPE on ONL and POS organization (*Figure S2a-S2h*). Both RO monoculture and RO-RPE co-culture showed the expected localization of POS (ROS and COS) markers, RCVRN (ROS and COS), RHO (ROS) and ML-OPSIN (COS) in the POS-like projections on the RO (*Figure S2c-S2h).* However, mislocalization of RCVRN (ROS and COS), RHO (ROS) and ML-OPSIN (COS) was also seen in the entirety of the three-to-five nuclei thick ONL (*Figure S2c-S2h)*. Furthermore, RCVRN^+^ (ROS and COS), RHO+ (ROS) and ML-OPSIN^+^ (COS) were also seen in the interior of the RO beyond the ONL (*Figure S2c-S2h).* Overall, the direct co-culture of RO and RPE did not improve photoreceptor cell maturation and localization in the RO.

Confocal imaging is routinely utilized to evaluate intact cells and tissues encapsulated in PEG hydrogel (*45–47*). Therefore, we primarily relied on confocal microscopy of tissue wholemounts for the analyses of the photoreceptors/RO in the PEG-HA RO and PEG-HA RO-RPE. Note that the encapsulation of RO in PEG-HA gels posed challenges for histological preparation due to the high water content of PEG-based gels (*45, 48, 49*). Although a few protocols have been optimized for preparation of cells/tissues encapsulated in PEG hydrogel for cryosectioning and transmission electron microscopy (TEM) (*45, 48, 49*), they were not suitable for the RO-RPE culture because the PEG-HA matrix was sandwiched between the RO and RPE monolayer. The RO and RPE physically separated and the fragile ROS and COS on the surface of RO and RPE were lost during tissue processing and sectioning (*Figure S3-S5*). Despite technical challenges, in a subset of experiments, we also utilized confocal microscopy and TEM to corroborate the structure of the photoreceptor and RPE monolayer and POS phagocytosis by RPE cells in tissue cryosections (*Figure S3-S5*).

Both the PEG-HA RO monoculture and PEG-HA RO-RPE assembloid model showed robust presence of RHO^+^ ROS and ML-OPSIN^+^ COS (*Figure 1g-1j and S3a-S3f*). However, similar to RO monocultures (in the absence of PEG-HA), mislocalization of ROS and COS was seen in the entirety of the ONL and interior of the PEG-HA RO (*Figure S3a-S3f)*. In contrast, in the PEG-HA RO-RPE, the localization of RHO and PNA-lectin and ML-OPSIN was predominantly restricted to the ROS and COS in the RO when imaged in the area of direct contact between RO and RPE monolayer (*Figure 1g-1i and S3a-S3f*). TEM and immunocytochemical analysis of the RO cross-section confirmed mature photoreceptors with presence of inner segments, outer limiting membrane (OLM), connecting cilium (CC) and outer segments (OS) at different stages of maturation including POS with well-organized stacks (*Figure S4a-S4k*).

Immunocytochemical analyses of RPE wholemounts and cryosections in the PEG-HA RO-RPE showed the expected localization of tight junction protein, ZO-1, RPE microvilli marker, EZR, apically-expressed POS engulfment receptor, Mer tyrosine kinase (MerTK) and RPE signature protein, RPE65 (*Figure S5a-S5d*). Furthermore, TEM and immunocytochemical analyses confirmed RPE ultrastructure and *in vivo*-like functional photoreceptor-RPE interaction, where POSs (ROS, COS) shed by rod and cone photoreceptor cells were phagocytosed by RPE cells in the 3D RO-RPE assembloid model (*Figure 1i-1k and S5e-S5h)*.

Morphological analyses of the ROS and COS ingested by the RPE in the 3D RO-RPE model showed *in vivo*-like ultrastructure with stacks of flat lobulated discs arranged in parallel stacks (*Figure 1k and S5g-S5h*). Furthermore, ROS and COS were observed at different stages of lysosomal degradation within the RPE monolayer of the 3D RO-RPE assembloid (*Figure 1k and S5g-S5h*). In addition, deposition of sulfated glycosaminoglycans (sGAGs) between day 1 and day 7 was consistent with the deposition of an IPM-like extracellular matrix in the 3D RO-RPE (*Figure 1l*). Finally, the assessment of biomechanical stiffness showed that Young’s elastic modulus of the PEG-HA hydrogel was comparable to the reported modulus of the human retina (∼10-20 kDa) (*50*) (*Figure 1m*).

To critically evaluate the role of matrix support in promoting the 3D RO-RPE model (*Figure 1*), we next compared the properties of the RO-RPE co-culture model in parallel experiments utilizing two distinct hyaluronic acid based MMP-degradable hydrogels, HyStem^®^ (https://advancedbiomatrix.com/hystem1.html) or the PEG-HA gels (*Figure S6*). PEG-HA gels are highly elastic with a linear stress-strain relationship (*Figure S6a*). In contrast, the HyStem^®^ gels were inelastic with poorly correlated stress-strain curve (*Figure S6a*). There was also less deposition of sGAGs in the RO layer in the HyStem^®^ RO-RPE when compared to the RO layer in the PEG-HA RO-RPE (*Figure S6b*). Furthermore, the distance between RO and RPE was greater in HyStem^®^ RO-RPE when compared to PEG-HA RO-RPE (*Figure S6c*).

RO encapsulated in both PEG-HA and HyStem^®^ gels showed robust presence of ROS and COS (*Figure 1g-1h and S6d-S6h*). However, POSs (ROS and COS) in the PEG-HA RO-RPE consistently showed the expected spatial apposition to the RPE which was in contrast to the HyStem^®^ RO-RPE, that displayed greater variability in POS (ROS and COS) juxtaposition with the RPE cells (*Figure 1g-1h and S6f-S6h*).

Overall, we have developed the PEG-HA hydrogel to provide optimal matrix support for co-culture of RO and RPE. The PEG-HA RO-RPE assembloid displays several properties of the *in vivo* retina, including *i)* biomechanical property of tissue elasticity, *ii)* sGAG deposition in the matrix, *iii)* structural spatial juxtaposition of POS (ROS and COS) to RPE monolayer, *iv)* POS (ROS and COS) ultrastructural morphology and maturation, and *v)* functional interaction of photoreceptor and RPE with phagocytosis and degradation of POS (ROS and COS).

### POS disorganization and POS loss in the RO-RPE model can be initiated solely by altered sphingolipid signaling due to RPE dysfunction

We next utilized the modular control of individual tissue layers (RO versus RPE) in the 3D RO-RPE model to investigate CLN3 disease mechanism. Note that clinical and histopathological analyses have shown that retina degeneration in CLN3 disease eyes initiates in the photoreceptor-RPE complex with disorganization and loss of POSs and proceeds inward into the retina (6, 44, 45). Furthermore, we have previously shown that RPE cells carrying CLN3 disease mutation (homozygous 966 bp deletion mutation spanning exon 7 and 8 in *CLN3; CLN3*^Δ*ex7-8*^) phagocytose POS less efficiently than control RPE cells (*10, 25*), but the pathological consequence of RPE dysfunction for POS loss in CLN3 disease is not known.

#### Spatially restricting CLN3 mutation to RPE cells in RO-RPE co-culture model is sufficient to instigate photoreceptor/retina degeneration

To determine the singular role of RPE dysfunction in promoting photoreceptor degeneration in CLN3 disease, control ROs (referred to as RO^control^) that showed robust presence of POSs (*Figure 2a*, *2b*) were co-cultured with isogenic control RPE (RPE^control^) versus *CLN3* mutant RPE (RPE*^CLN3^*) for a duration of 7 days (*Figure 2c*). A direct comparison of ROS and COS levels in the RPE layer of RO^control^ -RPE^control^ cultures versus RO^control^ -RPE*^CLN3^*cultures showed decreased number of ROS and COS within the RPE*^CLN3^*cells (*Figure 2d*). Furthermore, there were fewer RHO^+^ ROS and ML-OPSIN^+^ COS in the RO layer of RO^control^ -RPE*^CLN3^* cultures compared to RO^control^ -RPE^control^ cultures after 7 days of culture (*Figure 2e*). Also, when compared to RO^control^ - RPE^control^ cultures, ROS and COS in the RO layer of RO^control^ -RPE*^CLN3^* cultures were disorganized and did not display spatial juxtaposition with the RPE monolayer (*Figure 2e*).

**Figure 2.**
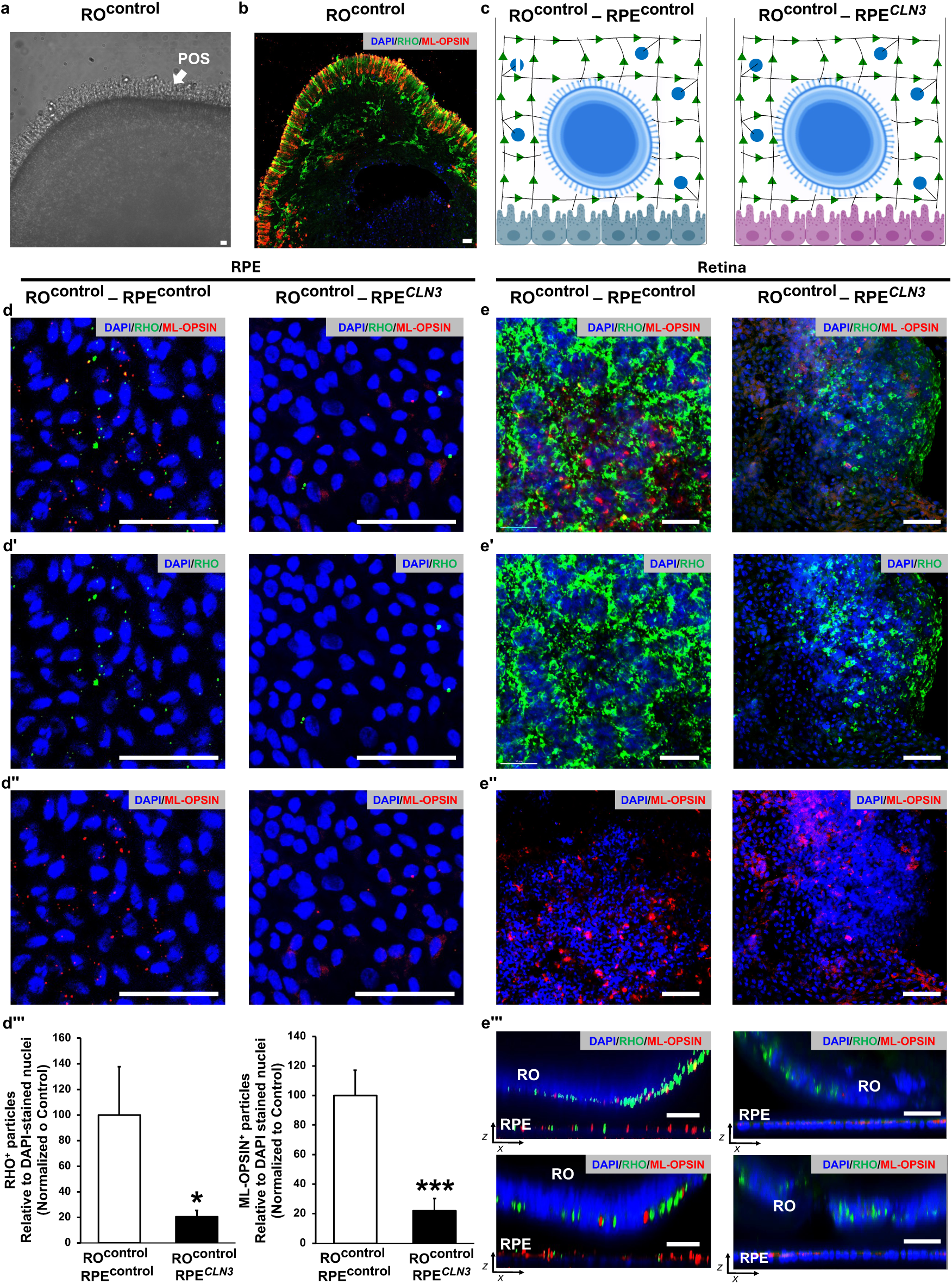
Spatially-restricting *CLN3* mutation to RPE cells is sufficient to cause POS disorganization and loss in the RO*^control^*-RPE*^CLN3^* assembloid model. **a, b)** Light microscopy (a) and immunofluorescence (b) images showing POSs (arrow, a) and RHO^+^ ROS (green, b) and ML-OPSIN^+^ COS (red, b) in the RO^control^ monoculture and prior to use in RO^control^-RPE*^CLN3^* assembloid culture. DAPI labeled nuclei are shown in panel b. Scale bar = 50 µm. **c**) Schematic representation showing the configuration of RO^control^-RPE^control^ versus RO^control^- RPE*^CLN3^* assembloid coculture where RO was derived from control hPSCs, but RPE monolayer was derived from control versus *CLN3* mutant hPSCs. **d)** Immunofluorescence images (d-d”) and quantitative analyses (d’’’) of RHO^+^ ROS (d-d’’’) and ML-OPSIN^+^ COS (d, d’’, d’’’) in the RPE monolayer RO^control^-RPE^control^ assembloid (left panels) and RO^control^-RPE*^CLN3^* (right panels) assembloids at day 7 of culture. Images are shown in planar view and cell nuclei are labeled with DAPI. Scale bar = 50 µm. * p<0.05, ***p < 0.005. **e**) Immunofluorescence images of RHO^+^ ROS (e-e’’’) and ML-OPSIN^+^ COS (e, e’’’) in day 7 RO^control^-RPE^control^ (left panel) and control RO^control^-RPE*^CLN3^* (right panel) assembloid cultures. Planar images of the RO layer are shown in panels e-e’’ and orthogonal xz views corresponding to two different planes are shown in panel e”’. Scale bar = 50 µm. Biological replicate n ≧ 3 for all Figure 2 experiments.

Overall, the comparison of RO^control^ -RPE^control^ and RO^control^ -RPE*^CLN3^* (*Figure 2*) shows that cell-autonomous *CLN3* mutation in RPE cells is independently sufficient to initiate disorganization of POS (ROS, COS) and POS loss in control RO. Furthermore, reduced levels of POS (ROS, COS) in RO^control^ -RPE*^CLN3^* compared to RO^control^ -RPE^control^ cultures is consistent with reduced uptake of POS by *CLN3* RPE that display reduced phagocytic function (*10, 25*).

#### Perturbation in sphingolipid signaling contributes to both reduced POS phagocytosis by CLN3 RPE cells and POS disorganization and loss in RO^control^ -RPE^CLN3^ cultures

We next utilized the hPSC-based molecular and cellular studies to determine “how” cell-autonomous *CLN3* mutation affects the photoreceptor-RPE interface and instigates POS disorganization and loss in the neural retina.

The RPE microvilli, which is densely packed with actin filaments and other cytoskeleton components, plays a key role in POS binding and consequently, POS phagocytosis. Our prior publication showed reduced density of RPE microvilli in CLN3 disease RPE cells compared to control RPE cells (*10*). In the RPE cell, EZR, a membrane-cytoskeleton linker protein, plays a critical role in apical microvilli formation (*51*). In particular, the membrane-bound, activated form of EZR is important for microvilli biogenesis (*52*). The comparison of EZR in the membrane versus the cytoplasmic fraction of isogenic RPE^control^ versus RPE*^CLN3^* cells showed reduced levels of EZR in the plasma membrane of RPE*^CLN3^* compared to plasma membrane of RPE^control^ (*Figure S7a, S7b*).

Sphingolipids regulate microvilli biogenesis, morphology, and stability in other epithelial cells (*53–55*). Furthermore, a critical sphingolipid signaling molecule, sphingosine-1-phosphate (S1P) has been shown to directly regulate EZR activation (*56*). *CLN3* deficient cells show decreased levels of N-acylsphingosine Aminohydrolase 1/Acid ceramidase (AC), a key regulator of sphingolipid metabolism (*57*). Analysis of *CLN3* expression in isogenic RPE^control^ versus RPE*^CLN3^* cells showed decreased *CLN3* transcript expression in RPE*^CLN3^* cells (*Figure S7c*). Consistently, AC levels were decreased in RPE*^CLN3^* cells compared to RPE^control^ cells (*Figure 3a*). In addition, the levels of AC-regulated sphingolipid, S1P, were reduced in RPE*^CLN3^* cells compared to RPE^control^ cells (*Figure 3b-3c*).

**Figure 3.**
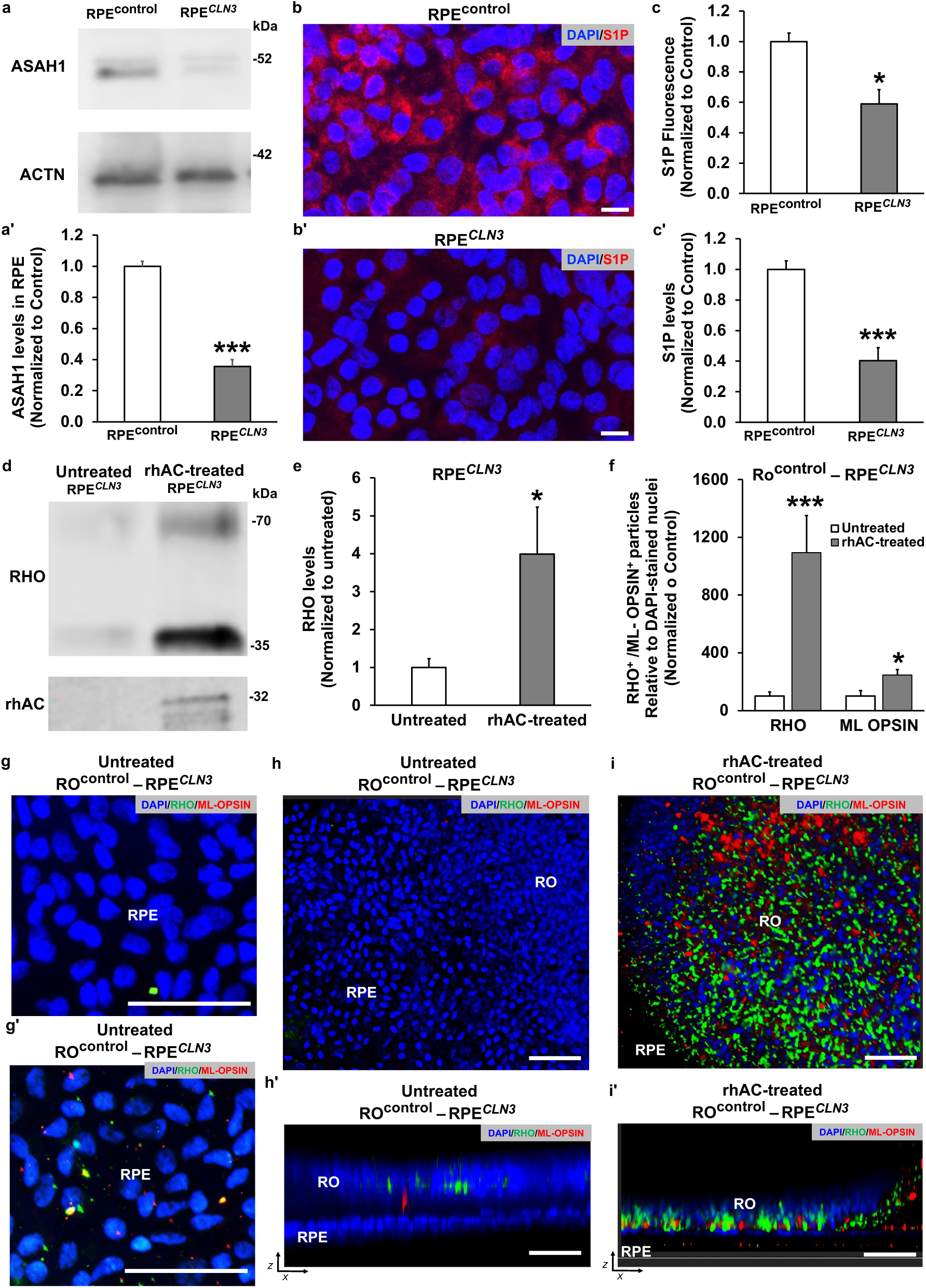
Recombinant acid ceramidase (rhAC) supplementation rescues POS (ROS and COS) loss in RO^control^ -RPE*^CLN3^* assembloid cultures. **a**) Western blot image (a) and quantitative analyses (a’) showing acid ceramidase AC (a, a’) and ACTN (a) levels in monocultures of RPE^control^ and RPE*^CLN3^* cells. ACTN served as the loading control and data are presented relative to ACTN and normalized to control sample. *** p<0.005. **b**) Immunofluorescence images showing sphingosine-1-phosphate (S1P) (red) localization in RPE^control^ (b) and RPE*^CLN3^* (b’) monocultures. DAPI^+^ cell nuclei are shown in blue. Scale bar = 50 µm. **c**) Quantitative analyses of S1P levels measured by immunofluorescence (c) and targeted lipidomics (c’) in RPE^control^ versus RPE*^CLN3^* monoculture. * p<0.05, *** p<0.01. **d, e)** Western blot image (d) and quantitative analyses (e) showing RHO and rhAC (d) and levels of RHO (e) in monocultures of untreated RPE*^CLN3^* and rhAC-treated (30µg/mL for 1h) RPE*^CLN3^* monocultures post ROS challenge (30 minutes, ∼20 ROS/RPE cells). Data in panel e is represented * p<0.05. **f, g**) Quantitative analyses (f) and immunofluorescence images (g, g’) illustrating the levels of RHO^+^ ROS and ML-OPSIN^+^ COS in the RPE monolayer of untreated RO^control^-RPE*^CLN3^* (f, g) and rhAC-treated (30µg/ml for 1h daily over 7 days) RO^control^-RPE*^CLN3^* (f, g’) assembloid cultures after 7 days of culture. Scale bar = 50 µm. * p<0.05, *** p<0.005. **h, i**) Immunofluorescence images showing the levels (h, i) and spatial organization (h’, i’) of RHO^+^ ROS and ML-OPSIN^+^ COS in the RO layer of untreated RO^control^-RPE*^CLN3^* (h, h’) and rhAC-treated (30µg/ml for 1h daily over 7 days) RO^control^-RPE*^CLN3^* (i, i’) assembloid cultures after 7 days of culture. Scale bar = 50 µm. Biological replicate n ≧ 3 for all Figure 3 experiments.

Therapeutically, recombinant human acid ceramidase (rhAC) treatment (30µg/mL for 1h) of RPE*^CLN3^* cells led to both increased S1P (*Figure S8a, S8b*) and plasma membrane-associated EZR (*Figure S8c, S8d*) compared to untreated RPE*^CLN3^* cells. Note that increased levels of intracellular S1P in rhAC treated cells is consistent with internalization bioactivity of rhAC in RPE*^CLN3^* cells. Supplementation of rhAC (30µg/mL for 1h) followed by POS/RHO phagocytosis assay (*10, 25*), confirmed uptake of rhAC by RPE cells and showed increase in the amount of POS/RHO uptake by rhAC treated RPE*^CLN3^* monocultures compared to untreated RPE*^CLN3^* monocultures (*Figure 3d, 3e*). Consistently, supplementation of rhAC (30µg/ml for 1h daily over 7 days) also led to increased levels of RHO^+^ ROS and ML-OPSIN^+^ COS in the RPE monolayer of rhAC-treated RO^control^ - RPE*^CLN3^* RPE assembloid cultures compared to untreated RO^control^ -RPE^control^ assembloid cultures (*Figure 3f, 3g*).

Also, compared to untreated RO^control^ -RPE*^CLN3^* cultures, rhAC-treated (30µg/ml for 1h daily over 7 days) RO^control^ -RPE*^CLN3^*cultures showed increased levels of ROS and COS in the RO layer of cultures (*Figure 3h, 3i and S8e, S8f*). Daily supplementation of rhAC also improved the spatial organization and juxtaposition of ROS and COS in the RO^control^ -RPE*^CLN3^* cultures compared to unsupplemented RO^control^ -RPE*^CLN3^* cultures (*Figure 3h, 3i*). The rhAC treatment for 7 days did not affect cell viability or epithelial integrity of the RPE*^CLN3^* and/or RO*^CLN3^* -RPE*^CLN3^*assembloid cultures (*Figure S9a-S9c*).

Altogether, these data show a direct role of AC-mediated sphingolipid signaling in *i)* impaired phagocytosis by RPE*^CLN3^*cells and *ii)* RPE-mediated photoreceptor degeneration in RO^control^ - RPE*^CLN3^* assembloid model.

### Perturbation in AC-S1P-EZR axis coincides with retina degeneration *in vivo*

Our *in vitro* studies showed a perturbation in AC-S1P-EZR axis in *CLN3* RPE cells (*Figure 3 and S7-S9*). To validate the impact of *CLN3* mutation on AC-S1P-EZR axis *in vivo*, we utilized homozygous *CLN3^Δex7-8^*miniswine (referred to as CLN3 miniswine), a large animal model of CLN3 disease that carries the orthologous mutation found in *CLN3* RPE cells and importantly displays CLN3 disease-associated retinal degeneration and vision loss (*58*).

#### Disorganization and loss of POS is seen in CLN3 miniswine retina at an early age

Longitudinal histological characterization of the wild-type (WT) and CLN3 miniswine retina was performed in the area centralis. We chose to focus on area centralis because it corresponds to the macula, the part of retina that shows the earliest pathological changes in CLN3 disease (*59, 60*). CLN3 miniswine retina showed POS (ROS and COS) disorganization at both 1-month and 6-months of age when compared to age-matched WT miniswine retina (*Figure 4a, 4b*). Note that prior studies that did not see overt retina pathology in CLN3 miniswine retina until 30-months of age (*25, 58*) did not consider the regional susceptibility of retina in diseases with macular involvement, like CLN3 disease.

**Figure 4.**
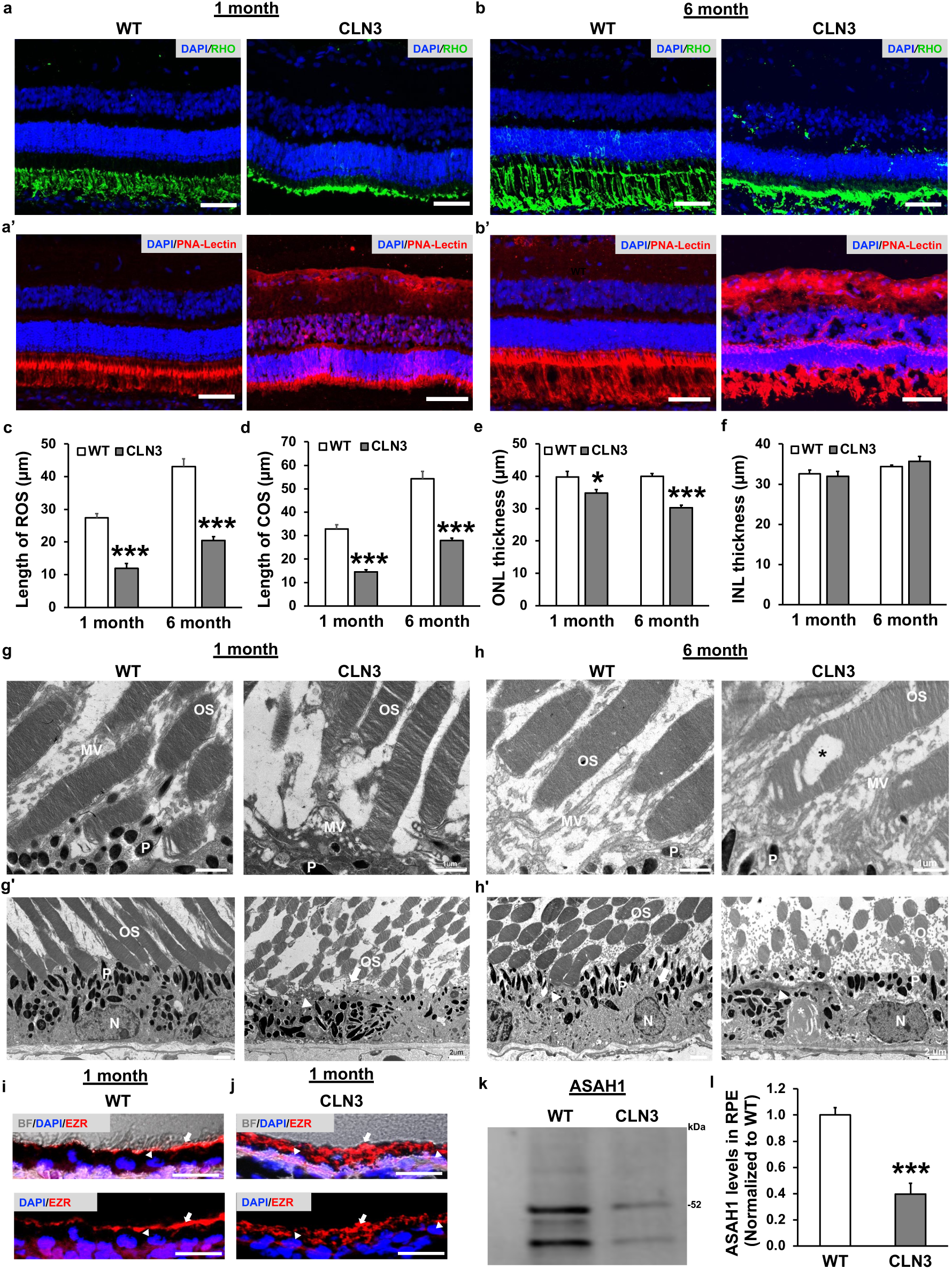
Longitudinal assessment of structural and molecular alterations in CLN3 miniswine model is consistent with RO^control^-RPE*^CLN3^* assembloid model. **a, b)** Immunofluorescence images showing RHO^+^ ROS (green; a, b), PNA-Lectin^+^ COS (red; a’, b’) in the wild-type (WT) and CLN3 miniswine retina cryosections at 1-month (a, a’) and 6-months (b, b’) of age. The ectopic localization of PNA-Lectin that is not restricted to COS is shown in a’, b’. Cell nuclei are labeled with DAPI (blue). Scale bar = 50 µm. **c, d**) Quantitative analyses showing the length of ROS (c) and COS (d) in wild-type (WT) and CLN3 miniswine retina at 1-month and 6-months of age. *** p<0.005. **e, f**) Quantitative analyses showing the thickness of the outer nuclear layer (ONL) (e) and inner nuclear layer (INL) (f) in wild-type (WT) and CLN3 miniswine retina at 1-month and 6-months of age. *p<0.05, *** p<0.005. **g, h)** Transmission electron microscopy images illustrating ultrastructure of POS, microvilli, and pigment granule in wild-type (WT) and CLN3 miniswine retina at 1-month (g, g’) and 6-months (h, h’) of age. A vacuole in 6-month CLN3 miniswine POS is shown with * (h). Disease associated fingerprint deposit is shown with * in CLN3 miniswine RPE (h’). OS, P, N, and MV denote POS, pigment, nuclei, and microvilli, respectively. Scale bars = 1 µm (g, h) and 2 µm (g’, h’). **i, j**) Brightfield and immunofluorescence images of retina sections showing the localization of EZR (red) in wild-type (WT) miniswine RPE (i) and CLN3 miniswine RPE (j). DAPI-stained RPE nuclei are shown in blue and indicated by white arrowheads. EZR is stained red and indicated by white arrow. Scale bar = 25 µm. **k, l**) Western blot images (k) and quantitative analyses (l) showing levels of AC (∼52 and 40 kDa) in wild-type (WT) versus CLN3 miniswine RPE at 1-month of age. Quantitative data for protein levels (l) are shown relative to total protein and normalized to wild-type RPE. *** p< 0.005. n ≧ 3 for 1-month and 6-month miniswine experiments in Figure 4.

Consistent with pathological manifestations spanning multiple retina cell layers in CLN3 disease, the localization of PNA-lectin that binds to a specific carbohydrate structure, galactosyl (β-1,3) N-acetylgalactosamine in COS was not restricted to POSs (COS) but was seen throughout the ONL and inner retina layers in CLN3 *miniswine* retina at both 1-month and 6-months of age (*Figure 4a, 4b*). This ectopic localization is perhaps not unexpected given that altered levels and sialylation of gangliosides, a key glycosphingolipid constituent of inner retina layers, has been documented in CLN3 disease (*62–64*). Furthermore, this is consistent with neuraminidase treatment removing sialic acid and thus causing altered PNA-lectin binding in the retina (*61*).

With regard to the outer retina pathology, apart from ROS and COS disorganization (*Figure 4a, 4b*), CLN3 miniswine retina also displayed decreased length of ROS and COS compared to age-matched wild-type miniswine retina at 1-month and 6-months of age (*Figure 4c, 4d*). Additionally, consistent with early photoreceptor degeneration, the thickness of the outer nuclear layer (ONL) was reduced in CLN3 miniswine retina in comparison to age-matched wild-type retina at both 1-month and 6-months of age (*Figure 4e, 4f*).

Ultrastructural analysis of the photoreceptor-RPE interface showed the typical, flat lobulated discs arranged in parallel stacks in wild-type and CLN3 miniswine retina at both 1-month and 6-months of age (*Figure 4g and 4h*). However, the 1-month- (*Figure 4g*) and 6-month-old (*Figure 4h*) CLN3 miniswine POS displayed apparent osmotic fragility, giving the appearance of swollen and disorganized disc membranes. Moreover, the OS structure of the 6-month-old CLN3 miniswine retina frequently displayed vacuoles that disrupted the disk membranes (*Figure 4h*). Lastly, CLN3 disease-associated fingerprint-like lipopigment was also observed in CLN3 miniswine RPE (*Figure 4h*).

In summary, the histological characterization of the CLN3 miniswine retina (*Figure 4a-4h*) was consistent with alterations at the photoreceptor-RPE interface occurring early in the disease.

#### Microvilli alterations and loss coincide with altered sphingolipid signaling in CLN3 miniswine RPE

The RPE monolayer in both wild-type and CLN3 miniswine retina showed presence of apical microvilli at 1-month-old (*Figure 4g*) and 6-months of age (*Figure 4h*). However, consistent with microvilli alterations in the RPE*^CLN3^* cells (*10*), the apical surface and microvilli of CLN3 miniswine RPE were highly convoluted and disorganized at both 1-month and 6-months of age (*Figure 4g, 4h*). Furthermore, compared to wild-type miniswine RPE, the levels of microvilli marker, EZR, were decreased in the CLN3 miniswine RPE at 1-month of age (*Figure S10a, S10b*). Also, consistent with EZR mislocalization to the cytoplasmic fraction in RPE*^CLN3^* monocultures (*Figure S7a, S7b*) and cytoskeletal defects in CLN3 disease (*20*), EZR did not display apical localization in CLN3 miniswine RPE monolayer at 1-month of age (*Figure 4i, 4j*). This was in contrast to the expected apical EZR localization in wild-type miniswine RPE at 1-month of age (*Figure 4i, 4j*).

Also, consistent with cytoskeletal defects in RPE cells, melanin pigment in 1-month- and 6-month-old CLN3 miniswine RPE were unevenly distributed in the cytoplasm and accumulated along the basal surface in some cells (*Figure 4g, 4h and S10c, S10d*). In contrast, and consistent with melanin localization in healthy RPE cells, the 1-month- (*Figure 4g, 4h, S10c and S10d*) and 6-month-old wild-type (*Figure 4h*) showed pigment granules distributed apically in the cytoplasm.

The evaluation of AC levels in wild-type and CLN3 miniswine RPE at 1-month of age also showed decreased AC in the CLN3 miniswine RPE (*Figure 4k, 4l*). Finally, consistent with AC-mediated alteration in sphingolipid signaling, total ceramide levels were increased in CLN3 miniswine RPE compared to wild-type miniswine RPE (*Figure S11a-S11c*).

Overall, molecular, and ultrastructural characterization of CLN3 miniswine retina is consistent with *CLN3* RPE and RO^control^ -RPE*^CLN3^* model and these data together suggest a role of altered RPE sphingolipid signaling contributes to POS loss and retina degeneration in CLN3 disease.

#### Histological evaluation of human donor eye is consistent with perturbation in AC-S1P-EZR axis

CLN3 disease donor RPE showed reduced levels of AC compared to age-matched control donor RPE (*Figure 5a, 5b*). Furthermore, S1P level was reduced in CLN3 disease donor RPE (*Figure 5c, 5d*). Also, consistent with perturbed AC-mediated sphingolipid signaling in CLN3 disease, elevated levels of ceramide were also observed in CLN3 disease donor retina/RPE, compared to age-matched control donor RPE (*Figure 5e, 5f*). Altered expression and localization of microvilli marker, EZR, was also seen in CLN3 disease RPE compared to age-matched control donor RPE (*Figure 5g, 5h*). In addition, CLN3 disease donor retina showed the expected loss of RHO^+^ ROS (*Figure 5i, 5j*) and extensive retina degeneration (*Figure 5e-5j*).

**Figure 5.**
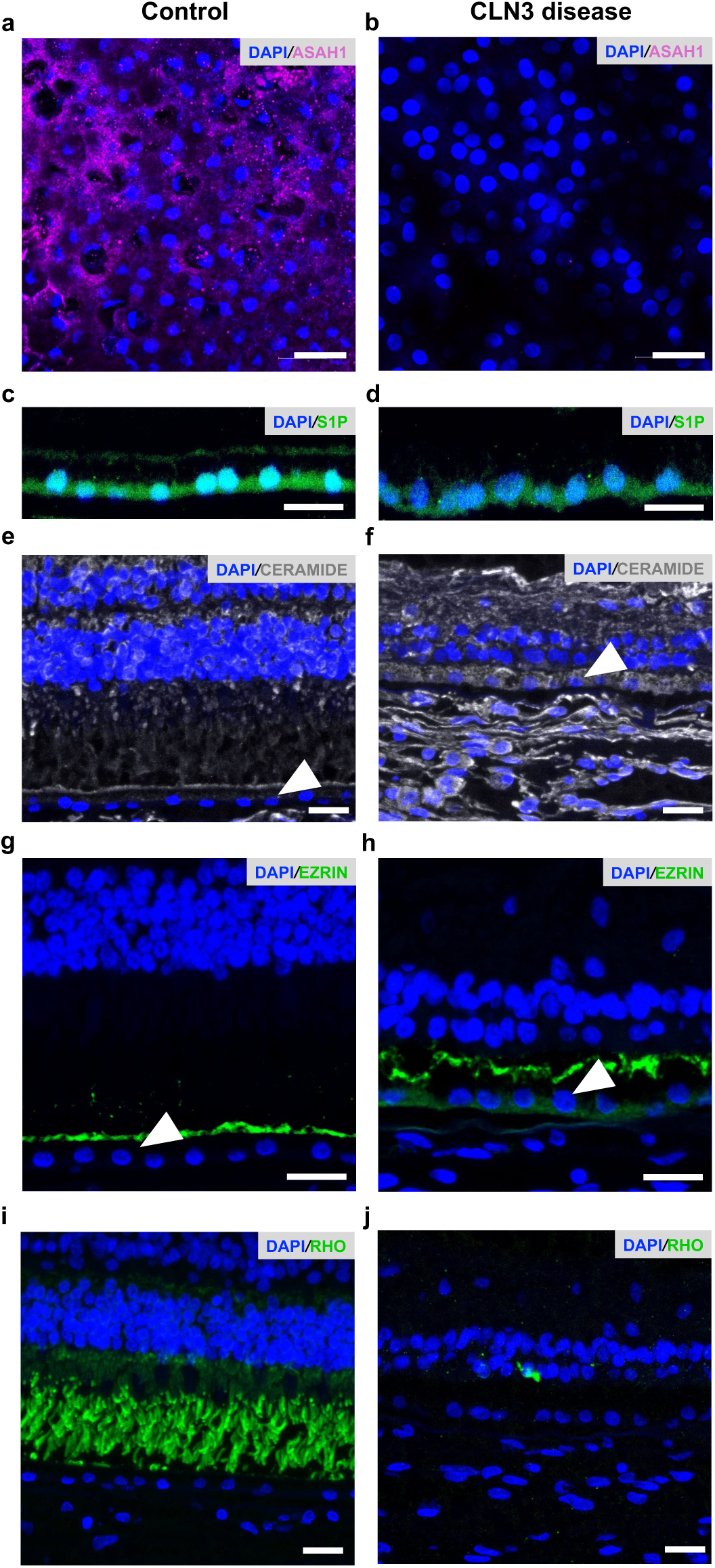
Structural and molecular alterations in CLN3 donor eye is consistent with control RO-CLN3 assembloid model. **a, b)** Immunofluorescence images showing AC (magenta) in RPE wholemount of an age-matched control donor eye (a) and CLN3 disease patient donor eye (b). Cell nuclei are labeled with DAPI (blue). Scale bar = 25 µm. **c, d)** Immunofluorescence images showing S1P (green) in RPE cryosections from an age-matched control donor eye (c) and CLN3 disease donor eye (d). DAPI labeled cell nuclei are shown in (blue). Scale bar = 25 µm. **e, f**) Immunofluorescence images showing total ceramide (white) and DAPI^+^ cell nuclei (blue) in retina cryosections from an age-matched control donor eye (e) and CLN3 disease donor eye (f). RPE nuclei is indicated with a white arrowhead. Scale bar = 25 µm. **g, h)** Immunofluorescence images showing microvilli protein, EZR (green) and cell nuclei (DAPI, blue) in retina cryosections from an age-matched control donor (g) and CLN3 disease donor (h). RPE nuclei is indicated with a white arrowhead. Scale bar = 25 µm. **i, j)** Immunofluorescence images showing RHO^+^ ROS (green) in cryosections of retina tissue from an age-matched control donor (i) and CLN3 disease donor (j). DAPI-labeled cell nuclei are shown in blue. Scale bar = 25 µm. n =1 for age-matched control donor eye and n=2 for age-matched CLN3 disease donor eye.

Overall, histological analyses of CLN3 donor eye were consistent with the RO^control^ -RPE*^CLN3^* assembloid model and supports a role of altered sphingolipid metabolism including perturbation of AC-S1P-EZR axis, microvilli disorganization and POS loss in CLN3 disease.

#### Longitudinal high resolution adaptive optics retinal imaging of human subjects with CLN3 disease is consistent with impaired POS phagocytosis preceding retina degeneration in CLN3 disease

Apart from histological analyses of animal/human donor retina progressing through the diseases, *in vivo* imaging of the living retina with an adaptive optics scanning light ophthalmoscope (AOSLO) provides valuable insight into cell-specific (e.g., individual photoreceptor versus RPE) changes in the retina (*65, 66*). Furthermore, adaptive optics fluorescence lifetime ophthalmoscopy (AOFLIO) allows evaluation of autofluorescence primarily from lipofuscin (POS digestion products) in the human photoreceptor-RPE complex (*67*).

We utilized AOSLO/AOFLIO imaging to compare RPE lipofuscin and photoreceptor cell health in normal subjects versus CLN3 disease patients. AOSLO/AOFLIO imaging was performed over regions of interest across the transition zone from atrophic to clinically intact retina in the CLN3 disease patient (*Figure 6*) (*68*). The same region of the retina was also imaged in a normal subject of comparable age (*Figure 6*). Images (1.5° square) with exposure times of 30-60 s have ∼0.5° of overlap between neighboring locations.

**Figure 6.**
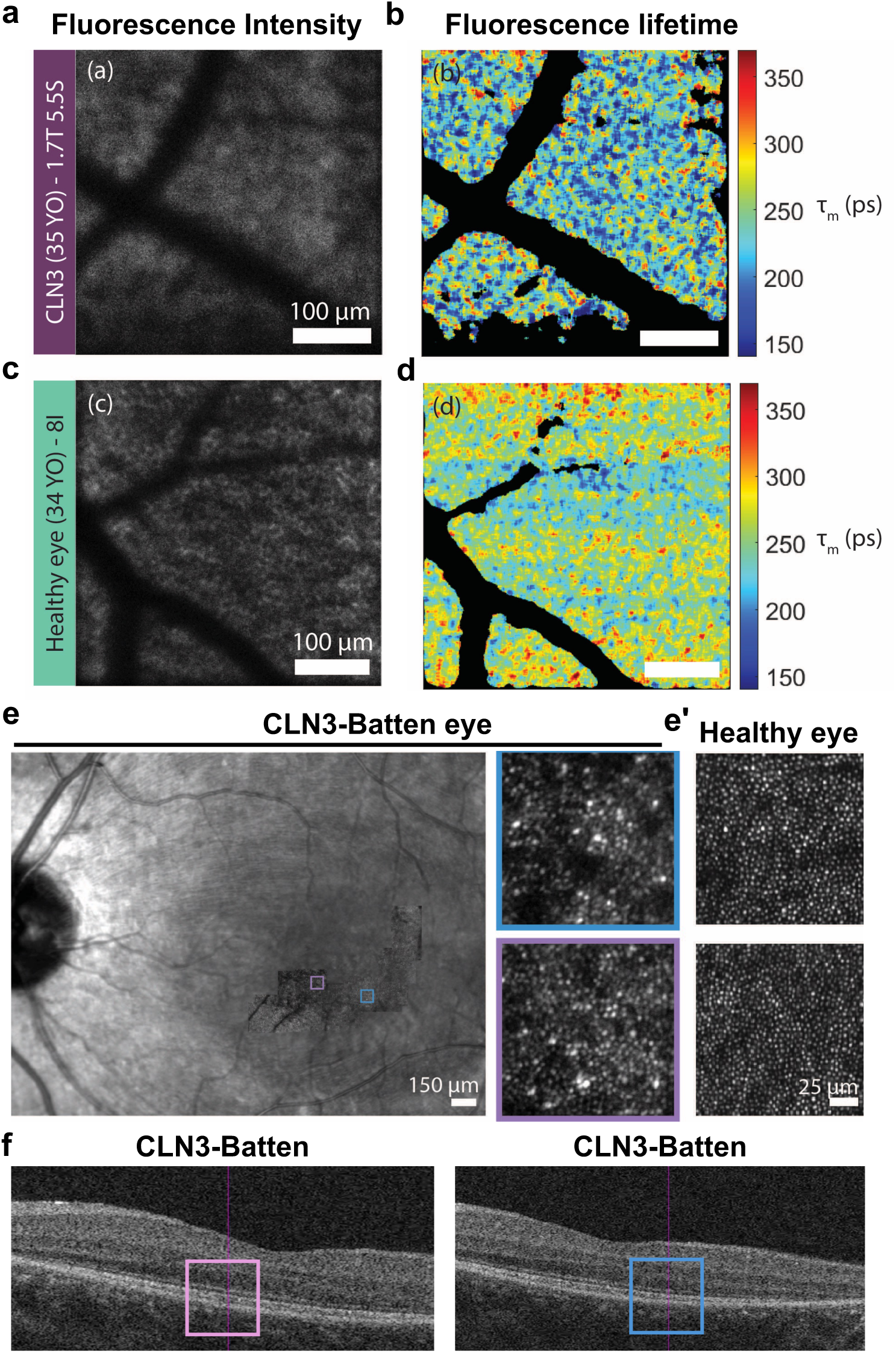
High resolution imaging supports decreased RPE lipofuscin and dysflective photoreceptors prior to retina degeneration in CLN3 disease. **a-d**) Adaptive optics fluorescence lifetime ophthalmoscopy (AOFLIO) measurement of fluorescence intensity (a, c) and fluorescence lifetime (b, d) in a CLN3 disease patient retina (a, b) and heathy retina (c, d). The measurement in the CLN3 retina were taken at 1.7 degrees temporal and 5.5 degrees superior in the middle of the atrophied region (a, b). The fluorescence intensity and fluorescence lifetime of an age-matched healthy retina was taken at a similar radial eccentricity of 8 degrees inferior (c, d). **e, f**) Adaptive optics scanning light ophthalmoscope (AOSLO) measurement of infrared reflectance (e) and spectral domain optical coherence tomography (SD-OCT) images (f) of a CLN3 disease patient retina. Higher magnification image of infrared reflectance of two eccentricity-matched locations (purple and blue square) are shown in CLN3 disease retina (e, right panel) and a healthy retina (e’). SD-OCT images in panel f indicate the location of magnified infrared reflectance (purple and blue square) in the CLN3 patient retina.

AOSLO/AOFLIO data support reduced lipofuscin levels in CLN3 disease RPE cells in living human patients (*Figure 6a-6d and S12a-S12d*). Furthermore, measurement of infrared reflectance taken on the AOSLO in eccentricity-matched locations showed more dysflective photoreceptors in CLN3 disease retina compared to normal healthy retina (*Figure 6e*). Consistent with dysflective cones preceding loss of POS and photoreceptor cell death, the spectral domain OCT (SD-OCT) imaging of the CLN3 disease patients in the area corresponding to dysflective cones in AOSLO imaging did not show evidence of overt retinal structural alterations including photoreceptor cell loss (*Figure 6f*).

Altogether, AOSLO/AOFLIO and OCT imaging support impaired POS phagocytosis and consequently POS disorganization and reduced RPE lipofuscin (partially-digested POS in RPE) in early-stage CLN3 disease prior to overt loss of photoreceptor cells.

#### Therapeutically targeting POS disorganization and loss in a stem cell model of CLN3 disease

Our prior results (*Figure 2, 3*) showed that primary RPE dysfunction leads to POS loss in the RO^control^ -RPE*^CLN3^* assembloid model. Furthermore, rhAC treatment rescued POS loss in RO^control^ -RPE*^CLN3^*assembloid model (*Figure 2*). To determine the therapeutic potential of rhAC for POS loss in CLN3 disease, we next investigated *i)* POS loss in the RO*^CLN3^* monoculture and RO^control^ - RPE*^CLN3^* culture and *ii)* effect of rhAC treatment on RO*^CLN3^*-RPE*^CLN3^* model.

#### *RO*^CLN3^ does not show loss of POSs (ROS, COS) in extended cultures

RO*^CLN3^* differentiated from a CLN3 disease hPSC line (harboring homozygous alleles of the common 966 bp deletion spanning exon 7 and 8) showed presence of VSX2^+^ bipolar cells and recoverin (RCVRN^+^) photoreceptors (*Figure S13a*). Furthermore, mature cultures of RO*^CLN3^* (>day 150 in culture) showed presence of POSs (ROS and COS) (*Figure 7a, 7b* and *S13a, S13b*). Note that POSs were consistently present in RO*^CLN3^* up to 1 year in culture (the longest point of investigation) (*Figure 7a, 7b*). Together, these data suggest that presence of disease causing *CLN3* mutation in the neural retina (RO) alone is not independently sufficient to promote POS (ROS and COS) loss in CLN3 disease.

**Figure 7.**
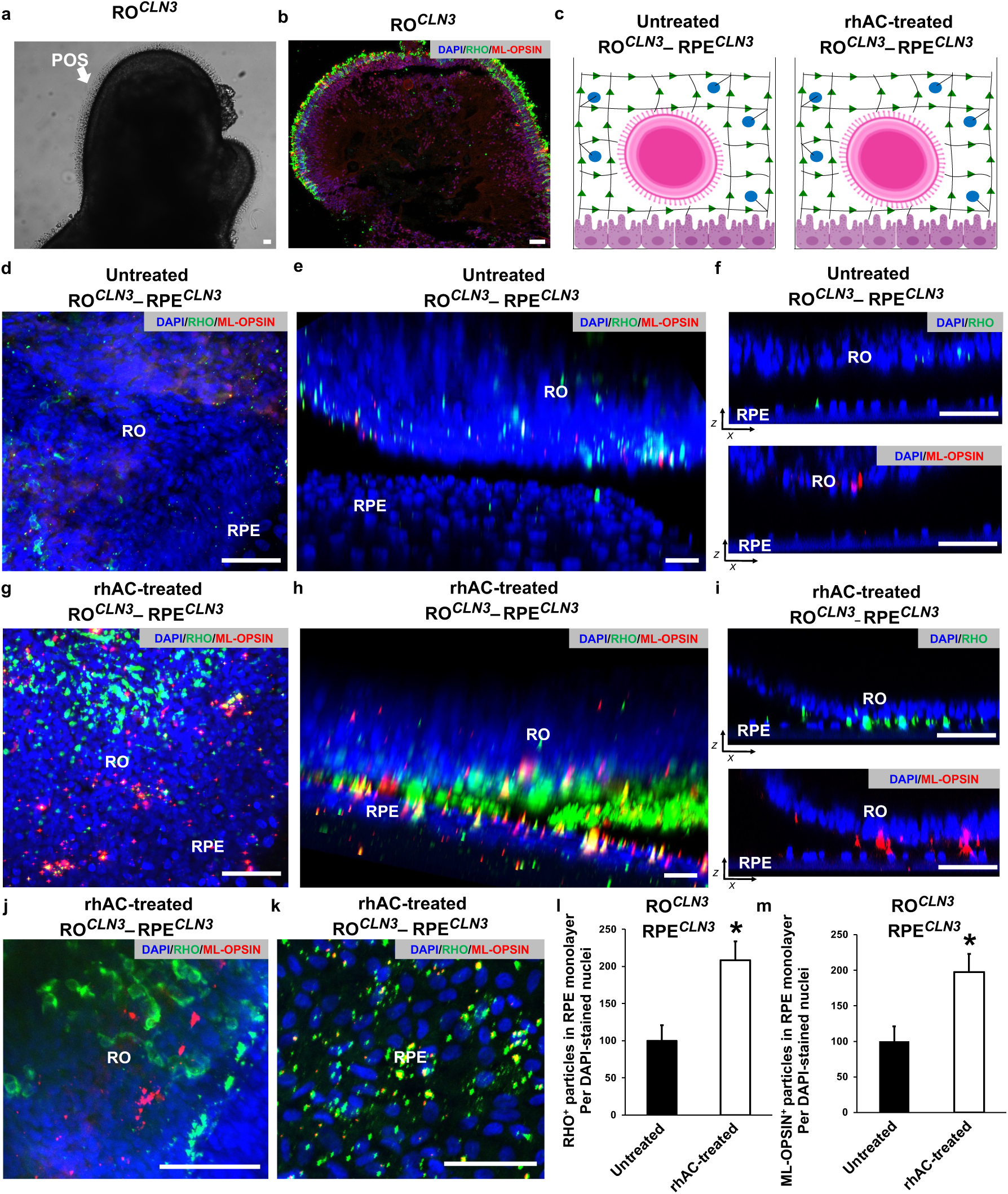
POS loss in the RO*CLN3*–RPE*CLN3* model can be rescued by rhAC supplementation. **a, b)** Light microscopy (a) and immunofluorescence (b) images showing POSs (arrow, a) and RHO^+^ ROS (green, b) and ML-OPSIN^+^ COS (red, b) in the RO*^CLN3^* monoculture and prior to use in RO*^CLN3^*–RPE*^CLN3^* assembloid. DAPI labeled nuclei (blue) are shown in panel b. Scale bar = 50 µm. **c**) Schematic representation showing the configuration of untreated and rhAC-treated RO*^CLN3^*– RPE*^CLN3^* cocultures where both RO and RPE were derived from CLN3 hPSCs. **d-f)** Immunofluorescence images in the planar view (d), 3D view (e), and orthogonal xz view (f) showing RHO^+^ ROS and ML-OPSIN^+^ COS in the RO and RPE monolayer of untreated RO*^CLN3^*– RPE*^CLN3^* assembloid at day 7 of culture. Cell nuclei (DAPI, blue). Scale bar = 50 µm. **g-i**) Immunofluorescence images of RHO^+^ ROS, ML-OPSIN^+^ COS, and DAPI^+^ cell nuclei in planar view (g), 3D view (h), and orthogonal xz view (i) in 7-day rhAC-treated (30µg/ml for 1h daily over 7 days) RO*^CLN3^*–RPE*^CLN3^* assembloid cultures. Scale bar = 50 µm. **j, k**) Immunofluorescence images of RHO^+^ ROS, ML-OPSIN^+^ COS, and DAPI ^+^ cell nuclei in the RO layer and RPE monolayer of rhAC-treated RO*^CLN3^*–RPE*^CLN3^* assembloid in planar view at day 7 of culture. Scale bar = 50 µm. **l, m**) Quantitative analyses of RHO^+^ ROS and ML-OPSIN^+^ COS in the RPE monolayer of untreated versus treated rhAC-treated RPE monolayer of the RO*^CLN3^*– RPE*^CLN3^* assembloid cultures. * p<0.05 Biological replicate n ≧ 3 for RO*^CLN3^* and RO*^CLN3^*– RPE*^CLN3^* cultures and n=2 independent experiments for rhAC treatments.

#### *RO*^CLN3^ shows loss of POS in the RO^CLN3^ -RPE^CLN3^ assembloid model

We next compared POS loss in isogenic RO^control^ -RPE^control^ versus RO*^CLN3^*-RPE*^CLN3^*assembloid cultures (*Figure S13c, S13d*). Similar to the RO in the RO^control^ -RPE*^CLN3^* cultures (*Figure 3e*), RO in the RO*^CLN3^*-RPE*^CLN3^* cultures showed loss of POSs (ROS and COS) compared to RO^control^-RPE^control^ cultures (Figure *S13c, S13d*). Furthermore, similar to the RPE in the RO^control^ -RPE*^CLN3^* cultures, RPE in the RO*^CLN3^*-RPE*^CLN3^* cultures also showed reduced levels of POSs (ROS and COS) compared to RO^control^ -RPE^control^ cultures (Figure *S13c, S13d*). Altogether, these data show that POS disorganization/loss in the RO and decreased POS in the RPE monolayer is similar in RO^control^ -RPE*^CLN3^* and RO*^CLN3^*-RPE*^CLN3^* assembloid cultures and is consistent with an independent role of primary RPE dysfunction in promoting photoreceptor degeneration in CLN3 disease.

#### rhAC rescues POS loss in the stem cell derived RO^CLN3^-RPE^CLN3^ model of CLN3 disease

Next, to investigate the therapeutic potential of rhAC for targeting photoreceptor degeneration in CLN3 disease, we examined the effect of rhAC treatment on POS organization and POS loss in RO*^CLN3^*-RPE*^CLN3^* model (*Figure 7c-7m*). rhAC supplementation (30 µg/ml for 1h daily over 7 days) of RO*^CLN3^-*RPE*^CLN3^* cultures led to both increased levels of POSs (ROS and COS) and improved spatial organization of POSs (ROS and COS) in the RO layer of the rhAC-treated RO*^CLN3^-*RPE*^CLN3^*cultures compared to parallel cultures of untreated RO*^CLN3^*-RPE*^CLN3^*cultures (*Figure 7c-7j*). Furthermore, rhAC supplementation daily for 7 days increased the number of POSs (ROS and COS) in both the RPE and RO of rhAC-treated RO*^CLN3^*-RPE*^CLN3^* cultures compared to parallel cultures of untreated RO*^CLN3^*-RPE*^CLN3^*cultures (*Figure 7k-7m and S13e, S13f*).

### Therapeutically targeting retina pathology in CLN3 miniswine eye

Our results thus far showed that rhAC supplementation can target POS phagocytosis defect in RPE*^CLN3^* cells (*Figure 3d, 3e*) and POS loss in hPSC-derived CLN3 “RO-RPE” model (*Figure.7c-7m and S13e, S13f*). Furthermore, longitudinal histological characterization of CLN3 miniswine eyes showed that decreased AC levels in CLN3 retina precede/coincide with RPE microvilli defects and POS disorganization and loss (*Figure. 4*). Therefore, to corroborate the therapeutic potential of rhAC supplementation *in vivo*, we utilized and evaluated the CLN3 miniswine model (*Figure 4*).

The longitudinal histological analysis of CLN3 miniswine retina (*Figure 4*) provided insights into an optimal timepoint for therapeutic intervention. Specifically, because the earliest timepoint of POS and RPE alterations were seen in CLN3 miniswine at ∼1 month (≥ 24 day of age), we chose to use day 20 CLN3 miniswine for rhAC supplementation *in vivo*. We next optimized the dose and route administration of rhAC for successfully targeting the retina/RPE monolayer in CLN3 disease. Prior studies have shown that systemically injected rhAC was bioavailable for at least 7 days (*69*). Based on the vitreal volume of miniswine eye (3-3.2 ml) (*70*), maximum allowed intravitreal injection volume in miniswine eyes (50 µl) and the target concentration of rhAC in the *in vitro* iRPE data (30 µg/ml), 48 µl volume of 1.92 mg/ml rhAC solution was intravitreally injected in the miniswine eye (*Figure 8a*). The contralateral eye was injected with the same volume of vehicle (48 µl 1X sterile PBS) (*Figure 8a*).

**Figure 8.**
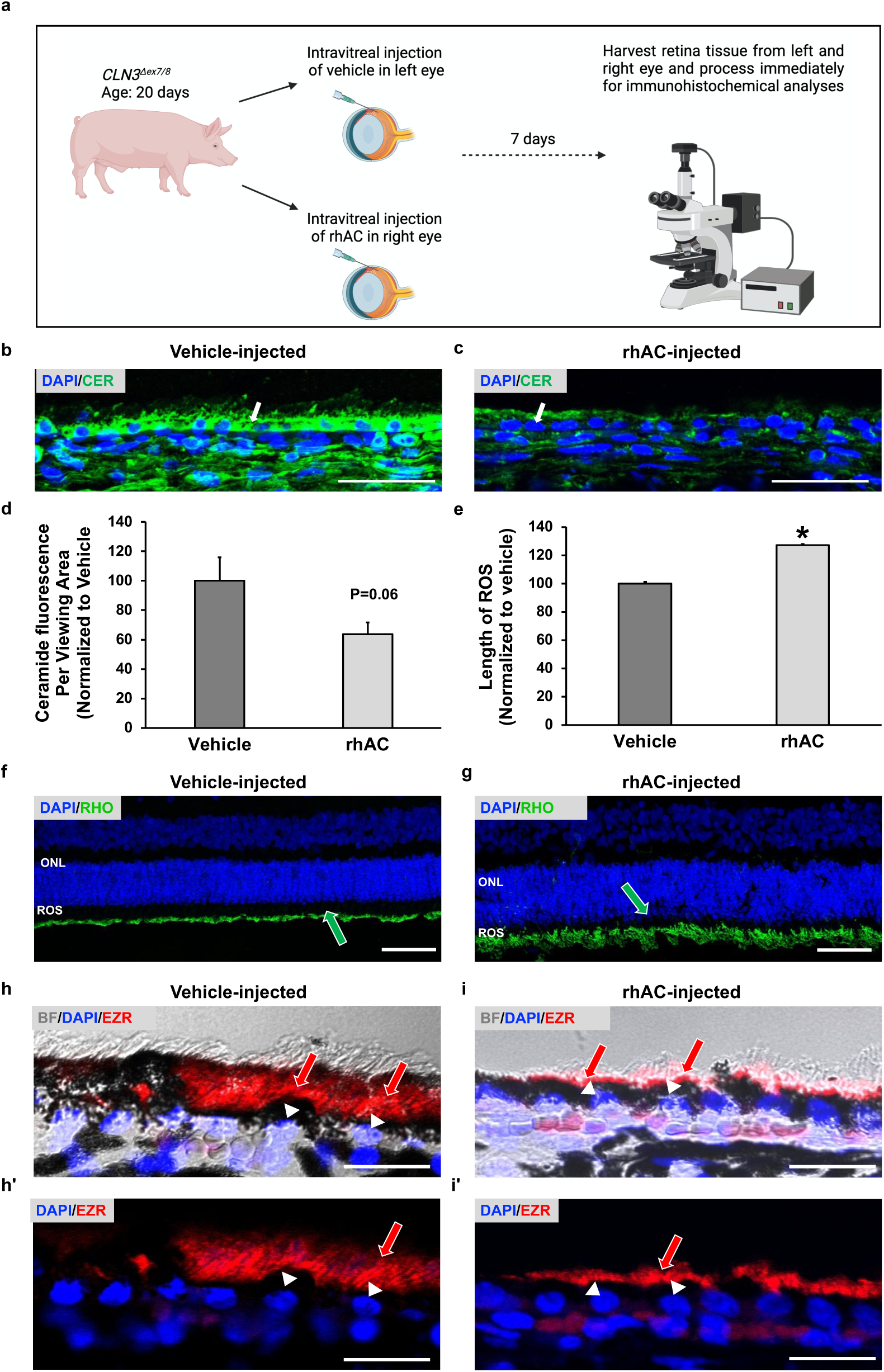
rhAC treatment targets retina pathology in the *CLN3* miniswine eye. **a)** Schematic representation of rhAC treatment in CLN3 miniswine eye. Equal volume of rhAC or vehicle was injected intravitreally in the right eye versus left eye of the CLN3 miniswine eye. Retina tissue from vehicle injected and rhAC injected CLN3 miniswine eye was harvested at day 7 of treatment and processed immediately for immunohistochemical analyses. **b-d)** Immunofluorescence images (b, c) and quantitative analyses showing total ceramide levels (green) in the RPE monolayer of the rhAC injected CLN3 miniswine RPE (b, d) compared to vehicle injected *CLN3* miniswine RPE (b, d) at day post-injection. Cell nuclei is labeled with DAPI and RPE monolayer is indicated with a white arrow in panel b, c. Scale bar = 50 µm. p=0.06. **e-g)** Quantitative analyses (e) and immunofluorescence images of retina sections showing RHO^+^ ROS (green; position indicated by green arrowhead) and DAPI ^+^ cell nuclei (blue) in vehicle injected (f) versus rhAC-injected (g) CLN3 miniswine eye at day 7 of treatment. Scale bar = 50 µm. *p<0.05. **h, i)** Immunofluorescence and brightfield images of retina sections at the same magnification showing EZR localization (red) and melanin pigment in vehicle injected (h, h’) versus rhAC-injected (i, i’) CLN3 miniswine eye at day 7 of treatment. Cell nuclei (DAPI, blue). EZR localization and position of DAPI-stained nuclei are also highlighted with a red arrow and white arrowhead respectively. Scale bar = 50 µm. Biological replicate n =3 for all experiments.

We next utilized immunohistochemical analyses of pig retina (area centralis) to evaluate the effect of intravitreal rhAC injection on AC-mediated sphingolipid metabolism in RPE cells and CLN3-associated early pathological changes observed in CLN3 miniswine retina at 1-month of age, namely, EZR and melanin pigment mislocalization in RPE cells and POS disorganization and loss in the photoreceptor layer (*Figure 8b-8l*). Consistent with bioavailability and bioactivity of intravitreally administered rhAC by RPE cells at least for 7 days, increased levels of AC and reduced levels of ceramide was seen in the RPE layer of rhAC-injected eye at day 7 post-injection when compared to the vehicle-injected eye at day 7 post-injection (*Figure 8b-8d and Figure S14a-S14d*). Furthermore, consistent with a global impact of rhAC on sphingolipid metabolism in the retina, qualitative immunofluorescence analyses showed that total ceramide was decreased and ASAH1 level was also increased in the neural retina and choroid in the rhAC-injected eye compared to the vehicle-injected eye at day 7 post-injection (*Figure 8b, 8c and Figure S14a-S14c*).

Therapeutically, localization of RHO was improved, and length of ROS was increased in rhAC-injected CLN3 miniswine retina compared to vehicle-injected CLN3 miniswine retina at day 7 post-injection (*Figure 8e-8g*). Qualitative immunohistochemical analyses also showed better PNA-LECTIN localization with less unexpected binding to the inner retina layers in rhAC-injected CLN3 miniswine retina (*Figure S14e-S14g*). Furthermore, improved EZR apical microvilli localization was seen in the RPE monolayer of the rhAC injected CLN3 miniswine eye at day 7 of treatment compared to vehicle injected CLN3 miniswine eye (*Figure 8h, 8i*). Also, directly linking AC-mediated sphingolipid signaling in cytoskeletal defects in CLN3 disease RPE cells (*Figure 4, S10*); qualitative evaluation showed predominantly apical localization of melanin pigment in rhAC-injected CLN3 miniswine RPE compared to vehicle-injected CLN3 miniswine RPE where melanin often spanned the entire width of the RPE monolayer (*Figure 8h, 8i*).

Finally, we evaluated the *in vivo* toxicity of rhAC in wild-type C57BL/6J mice. A dose equivalent to the rhAC concentration administered to CLN3 miniswine was injected in 1 eye of C57BL/6J mice. Equal volume of vehicle (1X PBS) was injected in the other eye. Both eyes were subjected to structural and functional evaluation using fundus imaging, fluorescein angiography (FA) and electroretinogram/ERG) 7 days-post injection (*Figure S14h-S14o*). Average a- and b-wave ERG amplitudes were similar between vehicle injected and rhAC injected mice 7 days post-injection (*Figure S14h-S14m*). Furthermore, no retinal anomalies were seen in fundus and FA examination in either the vehicle or rhAC injected mice at day 7 post-injection (*Figure S14n, S14o*).

Overall, disease modeling and therapeutic studies on the RO*^CLN3^*-RPE*^CLN3^*model and CLN3 miniswine eye supports a pathogenic role of reduced AC in CLN3 disease retina and identifies rhAC as potential therapeutic for CLN3 disease.

## DISCUSSION

In this study, we utilized a human retina cell model to study the pathobiology of CLN3 disease and identify a potential therapeutic. We chose to focus on the outer retina (photoreceptor-RPE complex) for studying CLN3 disease due to the importance of CLN3 function for outer retina homeostasis. *CLN3* is a ubiquitously expressed gene and there are currently >140 pathogenic *CLN3* variants that can lead to a spectrum of pathological manifestations (*71*). Specifically, C*LN3*-associated diseases include CLN3 disease, protracted CLN3 disease, and non-syndromic isolated retinal degeneration without neurologic involvement (*72–74*). The extensive characterization of retina in C*LN3*-associated disease including CLN3 disease, protracted CLN3 disease and CLN3-associated non-syndromic retinal degeneration has shown that all disease-causing *CLN3* mutations consistently affect the outer retina layers, the photoreceptor and RPE, and lead to photoreceptor cell death (*72–74*). In contrast, *CLN3* mutations variably impact other cell type(s) in the retina and brain and several studies have shown that inner retina degeneration and/or neurological impairment can be significantly delayed or absent in affected patients. Therefore, the outer retina (photoreceptor-RPE complex) provides an ideal tissue to investigate the role of *CLN3* gene/protein in normal and diseased physiology. Furthermore, CLN3 protein localizes to the lysosomes (*5–7*) and RPE cells have proven to be a suitable human cell model for studying lysosomal signaling and function in normal and diseased physiology. Therefore, we utilized RPE monocultures to investigate “how” CLN3 mutation impacts lysosomal sphingolipid signaling.

Our results show that the most common disease causing *CLN3* mutation (*CLN3*^Δ*ex7-8*^) leads to reduced levels of AC and perturbation of AC-mediated sphingolipid signaling contributes to cytoskeletal and phagocytosis defects in *CLN3* mutant RPE cells and POS disorganization loss in the photoreceptor layer of the *CLN3* mutant retina. Note that, although we utilized RPE cells for the investigation of AC levels in CLN3 disease, we expect reduced AC levels in other retinal and neuronal cell type(s) in CLN3 disease. Consistently, lysosomal proteome analysis of a cerebellar cell line derived from a *Cln3* knock-in mice with orthologous mutation (*Cln3*^Δ*ex7-8*^) showed decreased levels of AC (*57*). In addition, our data and published studies show ceramide accumulation in other cells, sera, retina, and brain of CLN3 disease patients and CLN3 disease mouse model (*23, 75–77*). We also expect that similar to the photoreceptor-RPE complex, reduced AC levels contribute to the broader pathological manifestations of CLN3 disease. Note that AC deficiency leads to “classical form of Farber disease”, another lysosomal storage disorder, that has a strong resemblance to CLN3 disease including retina degeneration, progressive neurological deterioration, and premature death (*78–82*).

Mechanistically, although we did not investigate “how” *CLN3*^Δ*ex7-8*^ mutation leads to reduced AC levels, CLN3 protein has been shown to impact lysosomal trafficking through Mannose-6-phosphate (M6P) receptors and this is a plausible explanation for reduced AC levels in the CLN3 disease cells. Therapeutically, our results shows that rhAC targeted several disease-associated phenotypes in the photoreceptor-RPE complex including altered sphingolipid metabolism (as measured by ceramide and S1P levels), pathological consequences of sphingolipid signaling on cytoskeletal organization (EZR and melanin localization) in RPE cells and POS disorganization and loss in the retina.

The evaluation of other cell layers in the wild-type versus CLN3 miniswine retina and vehicle-injected versus rhAC injected CLN3 miniswine eye supported the therapeutic potential of rhAC for targeting other pathological manifestations in CLN3 disease. For example, rhAC-injected CLN3 miniswine eye showed reduced ceramide and increased ASAH1 levels in the other retina cell layers including the photoreceptor and the choroid vasculature compared to vehicle-injected CLN3 eye. Similarly, rhAC targeted the inappropriate binding of PNA-LECTIN in the inner retina layers of CLN3 miniswine retina that potentially was a consequence of known alterations in sialylation and ganglioside accumulation in CLN3 disease (*62–64*). Note that directly linking AC deficiency to ganglioside accumulation in CLN3 disease, increased ganglioside accumulation has also been documented in “classical form of Farber disease” caused by AC deficiency (*83*).

Consistent with a role of AC-mediated sphingolipid signaling and altered lipid metabolism contributing to CLN3 disease, the therapeutic pipeline of CLN3 disease includes approaches to target glycosphiloglipid accumulation and neuroinflammation (Batten-/miglustat; NCT05174039) and PPAR-α activation to target lipid metabolism (PLX-200/gemfibrozil; NCT04637282). Notably, AC-mediated sphingolipid signaling modulates TRPML1 channel activation (*84, 85*), another therapeutic approach currently being investigated for CLN3 disease.

The investigation of the photoreceptor-RPE complex in CLN3 disease also enabled investigation of the precise pathogenic role of a non-neuronal cell (RPE) in independently promoting neuronal degeneration in CLN3 disease. Note that although we and others have previously shown altered pathology and function of non-neuronal cells in CLN3 disease (*10, 25–27*); the direct relevance of CLN3 function in non-neuronal cells for neurodegeneration in CLN3 disease was thus far lacking. By spatially restricting *CLN3* mutation to the RPE monolayer in the RO-RPE assembloid model, we were able to demonstrate that cell autonomous RPE dysfunction is sufficient to instigate POS disorganization and loss in CLN3 disease. Note that clinical and histopathological analyses have shown that retina degeneration in CLN3 disease eyes initiates at the POSs with POS disorganization and loss and proceeds inward into the retina (*9, 28, 29*). Therefore, it is plausible that RPE dysfunction is responsible for early retina degeneration in CLN3 disease and perhaps without RPE involvement, the neurons in the brain and retina would show similar susceptibility and time course of degeneration.

The RPE cells are specialized tissue-resident phagocytes in the eye that plays a key role in photoreceptor homeostasis (*86*). Furthermore, RPE cells are highly dependent on lysosomal function for their phagocytic activity and therefore it is not surprising that diseases affecting the lysosome (e.g., CLN3 disease) affect RPE cell health. Despite this, the role of RPE in lysosomal storage disorders including CLN3 disease has been under-investigated. Importantly, the direct involvement of RPE in promoting photoreceptor degeneration in CLN3 disease has direct relevance for several therapeutic approaches (e.g., gene therapy, ASO treatments) targeting retina degeneration in CLN3 disease.

From the perspective of broader understanding “how” lysosomal dysfunction including altered sphingolipid signaling affects RPE physiology has broader implications for several other retinal and neurodegenerative diseases including additional lysosomal storage disorders. Note that collectively lysosomal storage disorders are a common cause of progressive childhood neurodegeneration that affect 1 in ∼7500 live births (*87*) and vision loss due to retina degeneration is a common ocular manifestation of several lysosomal storage disorders (*88, 89*).

There are several limitations of our study. Although the 3D RO-RPE model provided evidence of *in vivo*-like POS maturation and RO-RPE interaction, the structural and functional evaluation of RO-RPE interaction was limited to POS abundance in the RO and RPE layer and the spatial organization of POS with respect to the RPE monolayer in the direct area of RO-RPE contact. The limited direct area of contact between the RO and RPE in the RO-RPE model was due to spherical versus planar geometry of the RO versus RPE monolayer. This was an important consideration for excluding other functional characteristics and global gene and protein expression studies in the evaluation of the 3D RO-RPE model. Current technical limitations like the detrimental effect of fixatives on tissue integrity, including separation of RO and RPE monolayer during tissue processing for electron microscopy and immunohistochemistry also limited the analysis of the comprehensive RO-RPE tissue. Other limitations of the study include lack of verification of the pathological RO-RPE interaction and rhAC testing in CLN3 patient-derived RO-RPE model. This was because the isogenic control and *CLN3* mutant lines provided a powerful approach to study the independent impact of the disease-causing *CLN3* mutation on photoreceptor-RPE interaction. In addition, although we used a large animal model (CLN3 miniswine) to perform proof-of-concept therapeutic studies testing impact of rhAC on early retina pathology in this model, we did not examine the long-term therapeutic consequences of rhAC in CLN3 miniswine model including evaluation of functional improvement of vision and broader pathological manifestations of CLN3 disease.

## MATERIALS AND METHODS

Detailed methods, including details of standard methodologies (e.g., cell culture, immunocytochemistry, and Western blotting) are provided in the Supplementary materials and methods.

### Procurement and Use of hESC Lines

H9 human embryonic stem cells (hESCs) were obtained from WiCell and used under approval from the Tasmanian Human Research Ethics Committee (#13502) and Institutional Regulatory Board (RSRB00056538) at the University of Rochester and conformed with the ethical norms and the declaration of Helsinki. CRISPR-Cas9-mediated editing of CLN3 was used to generate isogenic lines with and without the common 966bp deletion mutation spanning exon 7 and 8 (*25*). Isogenic control CLN3 mutant (referred to as CLN3) hESCs lines were used for differentiation into RPE and RO as previously described (*25*). Characterization of the pluripotency and karyotyping analyses of control and CLN3 hESC lines are previously described (*25*).

### Subject Details

Two CLN3 patients (ages 17 and 35 years) and four control subjects (ages 23, 29, 34 and 39 years) were used for imaging analyses following informed consent from the University of Rochester in accordance with an approved Institutional Regulatory Board protocol and conformed to the ethical norms and standards in the Declarations of Helsinki.

### Animals

Wild-type (WT) and transgenic (*CLN3^1′ex7-8^*) miniswine (*25, 58*) used in this study were obtained from Exemplar Genetics. All miniswine studies were performed following the approval of the Institutional Animal Care and Use Committees at Exemplar Genetics (Protocol # MRP2018-004). C57BL/6J mice used in this study were obtained from Jackson laboratory and all mouse studies were performed with the approval of University Committee on Animal Resources at the University of Rochester (Protocol # UCAR-2019-012).

### Experimental Setup and Statistical Analyses

Each individual experiment was carried out at least in triplicates. Data throughout the manuscript is presented as mean ± SEM. Statistical significance (* p < 0.05, ** p < 0.01, *** p < 0.005) was determined using unpaired Student’s t-test in Microsoft Excel.

## Supporting information

comprehesive supplementary file

## List of Supplementary Materials

Materials and Methods

Table S1 and S2

Figure. S1 to S14

## Acknowledgments

This work was supported by Exemplar Genetics staff who performed the intravitreal injections for the miniswine experiments. Furthermore, the electron microscopy facility at the University of Rochester supported the histological analysis of RPE cultures. Also, Dr. Robert Mullins at the University of Iowa provided expert insights into the histology data.

## Author contributions

Conceptualization and Methodology, R.S., D.S.W.B, V.B., and J.J.H; Investigation, J.H, N.F., S.D., J.A.H.T., A.C., L.K.K., A.K.S., Y.S., K.L, C.A.G., and A.H.; Visualization, V.B., C.A.G., D.S.W.B., and R.S.; Formal analysis, J.H., N.F., S.D., J.A.H.T., A.C., L.K.K., A.K.S., K.L., Y.S., D.S.W.B, R.L.T., and R.S.; Writing-original draft, R.S.; supported by J.H., N.F., S.D., and A.H.; Writing – Review & Editing, J.H., N.F., S.D., J.A.H.T, K.L., A.L.C., V.B., D.S.W.B, A.H., C.A.G., R.L.T., and R.S.; Resources, A.L.C., J.A.H; and E.S., Project Administration and Supervision, R.S., D.S.W.B. and J.J.H; Funding acquisition, R.S.

## Data and materials availability

Aside from use of established human pluripotent stem cell lines (hPSCs), and available transgenic miniswine model, there are no novel or unique resources that will be generated by the proposed research. In any case, our laboratory is committed to providing any assistance in reagent or protocol requests from other qualified investigators after publication of our experimental results. Information and requests for reagents may be directed to and will be fulfilled by the corresponding author, Dr. Ruchira Singh. Various hPSC lines have been established in this study from control subjects with no known history of retinal disease and will be readily shared with the scientific community upon request following the publication of experimental results by our laboratory. The subset of hPSC lines used in this study created by other investigators, would require permission and execution of a standard MTA by their institution for sharing of those lines, as would be required by our institution.

