## Supplementary material for "A 3D iPSC retina model reveals non-cell-autonomous and non-neuronal mechanism of photoreceptor degeneration in a lysosomal storage disorder": comprehesive supplementary file

### SUPPLEMENTARY MATERIALS

### MATERIALS AND METHODS

#### Differentiation of hESCs to retinal organoids (ROs)

hESC colonies were dissociated and cultured in suspension to generate embryoid bodies (EBs). On day 6, EBs were transferred to laminin-coated plates and maintained in neural induction medium (NIM, DMEM/F12 with 1% MEM-NEAA (Invitrogen, 11140-050), 1% GlutaMAX (Invitrogen, 35050-079), 1% N-2 Supplement (Invitrogen, 17502-010) and 2 µg/µL heparin (Sigma, H3149) and 1% (v/v) Penicillin and Streptomycin). On day 15, the cell growth medium was switched from NIM to 3D retinal differentiation medium (3D-RDM, 70% DMEM/30% F12 with 5% FBS (Invitrogen, A5669801), 1% MEM-NEAA, 1% GlutaMAX, 0.2 mM Taurine (Sigma, T8691), 0.5% Chemically Defined Lipids Concentrate (Invitrogen, 11905-031) and B-27 Supplement (Invitrogen, 17504-044) and 1% (v/v) Penicillin and Streptomycin). At ~day 20, RO-like structures were removed and transferred to polyHEMA (Sigma, P3932)-coated T25 flasks containing 3D-RDM. ROs were maintained in 3D-RDM. For tissue mimetic experiments, Stage 3 (~day 150) ROs with photoreceptor outer segments (POS) were selected for embedding in hydrogels.

#### Differentiation of hESCs to retinal pigment epithelium (RPE)

Differentiation of hESCs into RPE was conducted as previously described (*1-3*). Cell cultures were differentiated as described above. After removal of RO-like structures, the media was changed to retinal differentiation medium (RDM, 70% DMEM/30% F12 with B-27 Supplement (Invitrogen, 12587-010) and 1% (v/v) Penicillin and Streptomycin) and the remaining cells were allowed to grow as adherent cultures. RPE were dissected and dissociated with 0.05% Trypsin-EDTA and plated onto 24-well plates coated with laminin for 4-24 hours. RPE cells were thereafter cultured in RDM containing 2% FBS until the cells had formed a confluent monolayer. After reaching confluence, FBS was removed from the cell culture media and cells were maintained in RDM. For experiments, RPE from 24-well plates were dissociated with 0.05% Trypsin-EDTA and plated onto either 24-well plates again or 24-well ThinCerts™ inserts (6.5 mm, 0.4 µm pore, transparent, Greiner Bio-One, 662641) coated with laminin for 4-24 hours. RPE cells were thereafter cultured in RDM containing 2% FBS until the cells formed a confluent monolayer. On reaching confluence, FBS was removed from the cell culture media and cells were maintained in RDM. For tissue mimetic experiments specifically, RPE monolayers grown on transwell inserts were used for RO-embedded hydrogels.

**Synthesis of 8-arm Poly(ethylene glycol) (PEG)-amide Norbornene (PEG-NB)**

N,N'-dicyclohexylcarbodiimide (DCC, Sigma) coupling was used to synthesize norbornene-functionalized 8-arm PEG, as previously described (*4*). 20 g 8-arm 20 kDa PEG (JenKem Technology), pyridine (5 mol/mol PEG) and 4-dimethylaminopyridine (Fisher Scientific, 0.5 mol/mol PEG) were dissolved in 25 mL dichloromethane (DCM, Sigma) for 30 minutes. Separately, DCC (5 mol/mol PEG) and norbornene-2-carboxylate (10 mol/mol PEG, Alfa Aesar) were dissolved in 100 mL DCM for 30 minutes. Then, the PEG solution was added to the carbodiimide-activated norbornene, which was stirred overnight at room temperature. After filtration, PEG-NB was precipitated in cold diethyl ether and dialyzed against distilled, deionized water ($ddH_2O$, 1000 MWCO dialysis tubing, Spectrum Laboratories) for 48 hours and then frozen and lyophilized for three days. Norbornene-functionalization was measured by $^1$H-NMR [$CDCl_3$] to be 90% ($\delta$ = 3.5-3.8 of PEG ether protons compared to $\delta$ = 5.9-6.3 of norbornene vinyl protons).

N,N'-dicyclohexylcarbodiimide (DCC, Sigma) coupling was used to synthesize norbornene-functionalized 8-arm PEG, as previously described(*4*). 20 g 8-arm 20 kDa PEG (JenKem Technology), pyridine (5 mol/mol PEG) and 4-dimethylaminopyridine (Fisher Scientific, 0.5 mol/mol PEG) were dissolved in 25 mL dichloromethane (DCM, Sigma) for 30 minutes. Separately, DCC (5 mol/mol PEG) and norbornene-2-carboxylate (10 mol/mol PEG, Alfa Aesar) were dissolved in 100 mL DCM for 30 minutes. Then, the PEG solution was added to the carbodiimide-activated norbornene, which was stirred overnight at room temperature. After filtration, PEG-NB was precipitated in cold diethyl ether and dialyzed against distilled, deionized water ($ddH_2O$, 1000 MWCO dialysis tubing, Spectrum Laboratories) for 48 hours and then frozen and lyophilized for three days. Norbornene-functionalization was measured by $^1$H-NMR [$CDCl_3$] to be 90% ($\delta$ = 3.5-3.8 of PEG ether protons compared to $\delta$ = 5.9-6.3 of norbornene vinyl protons).

**MMP-degradable peptide synthesis:**

For in-house synthesis of MMP-degradable peptides, peptides were cleaved using 4 mL cocktail consisting of 0.5 mL triisopropylsilane, 0.5 mL $ddH_2O$, 0.5 mL 3,6-dioxa-1,8-octane dithiol, and 18.5 mL trifluoracetic acid for 30 min at 40$^o$C (CEM Razor - CEM, USA). Cleaved peptides were precipitated and washed three times in 100% diethyl ether, and vacuum-dried overnight (at least 18 h). Matrix-assisted desorption/ionization time of flight (**MALDI-TOF**) was used to verify the molecular mass of peptides(*5*). Peptides were purified by using XBridge® Peptide C18 column (19 mm x 150 mm, 5 µm particle size, 130Å pore size, 1/pKg from Waters Co, USA) using a gradient of acetonitrile and water over time (both solvents with 0.1% Trifluoroacetic acid – TFA) using peptide purification system Prodigy (CEM, USA). The samples were dissolved 10:1 (w:v – e.g., for each 100 mg peptide – 10 mL Act/$H_2O$/TFA) in acetonitrile:water with 0.1% TFA, each peptide had an optimal initial acetonitrile concentration. The volume injected during the purification process was locked in 5 mL per injection. MALDI was performed to verify the final product after purification. MALDI standards used covered a mass range ~1000-3200 Da using $\alpha$-Cyano-4-hydroxycinnamic acid as a matrix (Sigma Aldrich). The purification was validated using high-performance liquid chromatography (HPLC), and it is considered pure for this study when it is greater than 95% purity. The HPLC test was set up to 10 µL injection volume, and it was conducted using a mobile gradient phase consisting of gradient of acetonitrile:water with 0.05% TFA in a Kromasil C18 column (50 mm x 4.6 mm, 5 µm particle size, 100 Å pore size from Supelco, Bellefonte, PA) with a flow rate of 0.5 mL/min over 22 min. It used a fixed concentration of 2 mg/mL of peptide to perform the HPLC analysis. The column effluent was monitored with a variable wavelength UV−vis detector at 214 nm (Shimadzu Technologies). Peptides were freeze-dried (FreeZone 4.5 Liter -84C Benchtop Freeze Dryers – Labconco Co – USA) and stored at -80 °C until use. Stock solutions of peptides were stored at -80 °C until use.

**Immunocytochemistry**

Immunocytochemical analysis of RPE cell cultures was performed as previously described(*3*). Briefly, RPE on transwells were permeabilized and blocked in blocking buffer (10% normal donkey serum, 0.1% Triton-X-100) for 1 hour at room temperature. This was followed by overnight incubation in primary antibody in 0.5x blocking buffer at 4°C. The next day, RPE in transwells were washed with 0.05% Triton-X-100-PBS and subsequently incubated with host-specific Alexa Fluor-conjugated secondary antibody (1:500, Life Technologies) in 0.5X blocking buffer for 1 hour at room temperature. Following two 10 minutes washes with 0.05% Triton-X-

100-PBS, the transwell membranes were next incubated with Hoechst 33342 (H3570, Life Technologies) for 15 minutes to stain the nuclei, then washed in 1X PBS for 5 minutes. Subsequently, transwell membranes were cut out and mounted onto slides with ProLong Gold (P36930, Life Technologies) and imaged with a confocal microscope (LSM 510 META, Zeiss [Jena, Germany], or Eclipse Ti2, Nikon [Tokyo, Japan]). Primary antibodies used for immunocytochemical analysis included: S1P (Echelon Biosciences, Z-P300,1:50), ceramide (Enzo ALX-804-196-T050, 1:10), MERTK (1:100, Abcam), RPE 65 (1:100, GeneTex). Secondary antibodies used for immunocytochemical analysis included: donkey anti-mouse AF488 (1:500, Invitrogen, A21202), donkey anti-rabbit AF546 (1:500, Invitrogen, A10040).

For immunocytochemical analysis of RO and RO-RPE tissue sections, ROs were fixed in 4% PFA for 40 min at room temperature on a shaker, followed by washes with 1X PBS, 3 times. **Note:** For handling of ROs, wide orifice tips were used at all times. The fixed ROs were then incubated at room temperature with 15% sucrose-PBS for 60 mins or until the ROs sink to the bottom of the tube. This was followed by replacing the 15% sucrose-PBS solution with 30% sucrose-PBS solution and incubating the ROs at room temperature for 60 mins or until the ROs sink to the bottom of the tube. The ROs were then gently placed into a embedding molds carefully making sure there is no sucrose solution left in the mold. The mold was then filled with tissue freezing medium while the ROs were allowed to settle at the bottom of the embedding mold followed by freezing the RO blocks in a cryostat machine at -25$^{o}$C to -27$^{o}$C for at least 30 mins and store in -20$^{o}$C until ready for sectioning. Prior to immunostaining of the cryosectioned slides (14 µm thickness), a hydrophobic barrier was created around the sections to create ideal immunostaining conditions. The slides were warmed for 10 mins on a slide warmer followed by incubation with

1X PBS for 5 mins to remove the tissue freezing medium. This was followed by blocking in 1X blocking buffer at room temperature for 1 hour followed by incubation with primary antibodies solution made in 0.5X blocking buffer overnight at 4°C. The next day, the slides were incubated with 0.05% Triton-PBS solution for 10 minutes, 2 times, followed by incubation with host-specific Alexa Fluor-conjugated secondary antibody solutions (1:500, Life Technologies) diluted in 0.5X blocking buffer. Following this, the slides were again incubated with incubated with 0.05% Triton-PBS solution for 10 minutes, 2 times at room temperature followed by Hoechst (1:1000, Thermo Fisher, MA) at room temperature for 30 minutes and then mounted with ProLong Gold (P36930, Life Technologies) and imaged with a confocal microscope (LSM 510 META, Zeiss [Jena, Germany], or Eclipse Ti2, Nikon [Tokyo, Japan]). Primary antibodies used for immunocytochemical analysis of ROs included: RCVRN (1: 1000, Proteintech), VSX2 (1:200, Exalpha), Alexa Fluor® 488 phalloidin (1:100, ThermoFisher Scientific), CRALBP (1:200, Abcam), TUBB3 (1:200, Biolegend). Secondary antibodies used for immunocytochemical analysis included: donkey anti-mouse AF488 (1:500, Invitrogen, A21202), donkey anti-mouse AF546 (1:500, Invitrogen, A10036), donkey anti-rabbit AF488 (1:500, Invitrogen, A21206), donkey anti-rabbit AF546 (1:500, Invitrogen, A10040), donkey anti-rabbit AF647 (1:500, Invitrogen, A31573), donkey anti-sheep AF633 (1:500, Invitrogen, A21100).

For wholemount immunocytochemical analysis of PEG-HA RO and PEG-HA RO-RPE, the cultures were fixed in 4% paraformaldehyde for 30 minutes. Subsequently, fixed hydrogel samples were permeabilized and blocked in blocking buffer for 1 hour two times. Next the hydrogel samples were incubated with primary antibodies in 0.5X blocking buffer overnight at 4°C. The next day, following 3 hours of washes in 1X PBS, hydrogel samples were incubated in host-specific Alexa Fluor-conjugated secondary antibody (1:500, Life Technologies) in 0.5X blocking buffer overnight at 4°C. Subsequently, following 3 hours of washes in 1X PBS, hydrogel samples were incubated in anti-lectin PNA-AF647 (1:50, Invitrogen) and Hoechst (1:1000, Thermo Fisher, MA) at room temperature for 30 minutes. The hydrogels were mounted with Aqua-Poly/Mount (Polysciences, Warrington, PA) on depression slides (Fisher Scientific). All images were captured on a confocal microscope (Eclipse Ti2, Nikon, Tokyo, Japan). Primary antibodies used for immunocytochemical analysis included: RHO (1:100, Millipore, MABN15), M/L Opsin (1:100, Millipore, AB5405), CASP3 (1:100, Cell Signaling Technologies, 9602S). Secondary antibodies used for immunocytochemical analysis included: donkey anti-mouse AF488 (1:500, Invitrogen, A21202), donkey anti-rabbit AF546 (1:500, Invitrogen, A10040), donkey anti-rabbit AF647 (1:500, Invitrogen, A31573). List of primary antibodies used for immunocytochemistry are also provided in *Table S2*.

**Western Blotting**

Cells were lysed in RIPA buffer (89900, Thermo Fisher Scientific, MA) containing protease inhibitor (P2714, Sigma-Aldrich, St. Louis, MO). 30 μg of protein lysates were resolved on 4-20% gels by SDS-PAGE. Following SDS-PAGE, the proteins were transferred onto PVDF membranes, and the membranes were blocked for one hour at room temperature in either 5% non-fat dry milk (NFDM) in PBS or commercially bought blocking solution (927-65001, LI-COR Biosciences, NE). The membranes were then incubated overnight with anti-Rhodopsin (Millipore MABN15), anti-Ezrin (Cell Signaling Technologies, 3145S), anti-Caveolin 1 (Cell Signaling 3267), anti-Phosphoezrin (Invitrogen PA5-37763), anti-actin (Santa Cruz, SC-47778) antibodies in Licor blocking buffer at 4°C. The following days, membranes were washed with 1X PBS-T (0.1%

Tween-20), following incubation with host-specific HRP- or IR Dye-conjugated secondary antibody (1:10000) at room temperature for 1 hour. The membrane was subsequently washed thrice in 1X PBS-T and analyzed using Azure C-series chemiluminescence documentation system (Azure Biosystems, Dublin, CA, USA). ImageStudio (LI-COR Biosciences, NE) and Microsoft Excel software were utilized for quantification of bands. In a subset of experiments, subcellular fractionation of RPE cells was performed to collect plasma membrane and cytosolic fractions using the commercially available kit (BSP002 BioBasic In, ON). Following manufacturer's instructions, each fraction was collected and stored at -80°C until use for Western blot analysis. Primary antibodies used for Western blot analysis included: EZR (1:1000), CAV1 (1:750), ASAH1 (1:500), ACTN (1:500), RHO (1:500). List of primary antibodies used for western blotting are also provided in *Table S2*.

**Miniswine pig retina tissue collection and immunohistochemistry analyses**

***Miniswine pig RPE wholemount staining***. Miniswine pig eyes were harvested at 1 and 6 months of age. Eyes were injected with 4% PFA following enucleation for 1 hour prior to dissection to remove the anterior portion and vitreous. The remaining retina/RPE eyecup was submerged in 4% PFA for a fixation of at least 5 hours at 4°C. Following two 1X PBS washes, samples were prepared for wholemount staining by carefully removing the RPE/choriocapillaris from the sclera. Wholemount samples were permeabilized/blocked in blocking buffer (1% normal donkey serum (ImmunoReagents, Inc.), 0.2% Triton-X-100, 0.2% Tween-20) for 1 hour at room temperature. Primary antibody incubation was done in 0.5X blocking buffer at 4°C overnight. Samples were then washed 2-4 times with 0.05% Triton-X-100 in PBS and then incubated with host-specific Alexa Fluor™ secondary antibody (1:500, Life Technologies) in 0.5X blocking buffer for 1 hour at room temperature. The wholemount samples were washed again 2-4 times with 0.05% Triton-X-100 in PBS and incubated with DAPI dye (Hoechst 33342; Life Technologies) for 15-30 minutes at room temperature. Samples were briefly incubated with TrueBlack® (Biotium #23007) following manufacturer's instructions to quench autofluorescence signal. Samples were mounted onto slides with Prolong Gold (Life Technologies) and cover slipped.

***Immunohistochemistry staining on miniswine pig retina tissue.*** Miniswine pig eyes were harvested at 1 and 6 months of age. Immediately after enucleation, eyes were placed in 4% PFA. The anterior segment was carefully removed. Vitreous and lens were removed with extra effort to keep retina intact and attached. The posterior eyecup was placed back into 4% PFA for 4 hours at room temperature. This was followed by incubations with serial sucrose solutions: 10% 1h, 20% 1h, 30% overnight and then embedding prior to sectioning at 14 µm thickness. The sections were hydrated in 1X PBS followed by immunostaining as described above. Primary antibodies used for immunocytochemical analysis included: RHO (1:100, Millipore, MABN15) and Alexa Fluor™ 647 Conjugate PNA lectin (1:50, Invitrogen, L32460). Secondary antibodies used for immunocytochemical analysis included: donkey anti-mouse AF488 (1:500, Invitrogen, A21202). List of primary antibodies used for staining of pig retina tissue are provided in *Table S2*.

**Wholemount staining of human donor retina/RPE tissue**. Whole eye tissue sample from a 22-year-old donor with genotype-phenotype confirmed CLN3 disease (#591; time of enucleation: 4 h) and from a healthy adult without any history of retinal disease (time of enucleation: 10.2 h) was acquired from Dr. Vera Bonilha at Cleveland Clinic. Wholemount RPE samples were first treated with melanin bleach kit (Polysciences, 24883-1) following the manufacturer's instructions.

Following the melanin removal, immunostaining was performed on RPE wholemounts as described above. Human RPE wholemounts were stained with ASAH1 antibody (Millipore Sigma, ABN468, 1:100) and Alexa Fluor™ 633 secondary antibody (1:500, Life Technologies) Post-staining, the samples were incubated in TrueBlack® Lipofuscin Autofluorescence Quencher (Biotium, CA, USA, cat# 23007) following the company's instructions. Samples were then rinsed with 1X PBS and coverslipped with ProLong Gold mounting medium prior to imaging with Nikon Ti2 confocal microscope. Secondary antibodies used for immunocytochemical analysis included: donkey anti-sheep AF633 (1:500, Invitrogen, A21100). Primary antibodies used for staining of human donor retina/RPE tissue are listed in *Table S2*.

**Characterization of the PEG hydrogel tissue mimetic**

PEG-NB (7.5 wt%), MMP sensitive crosslinker peptide (10.8 mM, resulting in 80% crosslinking with PEG-NB) and 100 nM HA-thiol (1000 kDa, 5% functionalization, Creative PEGWorks, NC) were dissolved in Dulbecco's phosphate buffered saline (DPBS). 0.05 wt% lithium phenyl-2,4,6-trimethylbenzoylphosphinate (LAP), synthesized as described (Fairbanks et al., 2009), was then added to the hydrogel precursor solution. LAP is synthesized in-house using established methods(*6*). Acellular hydrogels (~Ø6 x 2 mm, 50 µL/gel) were photopolymerized using 365 nm light at 5 $mW/cm^2$ for 3 minutes. After incubation in DPBS overnight, the compressive modulus of the hydrogels was measured to be 14.5 ± 0.8 kPa using an MTS QT/5 (2 N load cell). HyStem® (Advanced Biomatrix, Carlsbad, CA, USA, GS312F) hydrogels were synthesized according to the manufacturer's instructions. Briefly, Glycosil (HA-thiol), Gelin S (gelatin-thiol) and Extralink (PEG-diacrylate) were resuspended in their respective buffers and combined in a 2:2:1 (Glycosil:Gelin S:Extralink) ratio. Acellular hydrogels (~Ø6 x 2 mm, 50 µL/gel) were polymerized at room temperature for 90 minutes. After incubation in DPBS overnight, the compressive modulus of the hydrogels was measured to be 5.1 ± 0.4 kPa using an MTS QT/5 (2 N load cell).

To entrap ROs within hydrogels, 65 μL of either PEG-HA or HyStem® hydrogel precursor solution, described above, was poured over 1-3 whole Stage 3 (presence of POS, ~>150 days) ROs, which were briefly resuspended in the solution using a wide-bore pipet tip and subsequently photopolymerized in a 24-well ThinCerts™ insert and cultured using 3D-RDM, which was exchanged every 2 days. For RO-RPE cocultures, ~50,000 RPE were first cultured to maturity on the transwell membrane surface (~30 days), and then RO-encapsulated hydrogels were photopolymerized on top of the RPE monolayer. To provide optimal culture conditions to both RO and RPE layer in the transwell model; RO monocultures were solely supplemented with media used to promote RO maturation whereas RO-RPE co-cultures were supplemented apically with media used to promote RO maturation and basally with media used to support RPE (*7, 8*).

**1,9-dimethylmethylene blue (DMMB) assay to determine sulfated glycosaminoglycan (sGAG) content**

Control RO was encapsulated in hydrogel and cocultured with RPE as described above for 24 hours or 7 days. After the coculture duration, the hydrogel-encapsulated RO was briefly dried and collected in a 1.5 mL microcentrifuge tube and stored at -20°C until ready for use. PBE buffer (pH 6.5) was prepared by dissolving 100 mM sodium phosphate dibasic ($Na_2HPO_4$, Sigma, 567547) and 10 mM ethylenediaminetetraacetic acid disodium salt (EDTA-$Na_2$, Sigma, 324503) in dd$H_2O$ adjusting the pH to 6.5. A papain digestion solution was then prepared by dissolving 1.75 mg/mL L-cysteine (Sigma, C5360) and 0.5% (v/v) papain (Worthington Biochemical, Lakewood, NJ, USA, LK003176) in PBE buffer. 300 μL papain was added to each hydrogel-encapsulated RO and homogenized using a tissue homogenizer. The total volume of the papain-hydrogel-RO suspension was subsequently brought to 500 μL using papain. Papain-hydrogel-RO suspensions were incubated at 60°C for 16 hours. Digested solutions were added at 1:1 to DMMB (Amsbio, Cambridge, MA, USA, 280560) in duplicate wells of an optically clear 96-well plate and allowed to incubate at room temperature for 5 minutes. Absorbance readings were taken at 522 nm wavelength using a plate reader (Varioskan Flash, Thermoscientific) and compared to a standard curve derived from chondroitin sulfate (amsbio, 280560).

**POS phagocytosis assay**

POS phagocytosis assay was performed *in vitro* as previously described(*1, 2*). Bovine POS was obtained commercially from InVision BioResources (Cat. #98740, Seattle, WA, USA). POS uptake by RPE cells was evaluated by feeding mature monolayer of RPE cells in culture with unlabeled POS (approximately 20-40 POS/RPE cell) for 30 min at 37°C with 10% FBS media supplementation. Thereafter, to remove POS (unbound) on the RPE cell surface, RPE cells were thoroughly washed with 1x PBS. Subsequently, RPE cells were harvested for Western blotting.

**Adaptive optics retinal imaging of human subjects**

To compare *in vivo* photoreceptor quality and RPE fluorophore composition, the retinas of 2 CLN3 (17 and 35 years old) and 4 young (23, 29, 34, and 39 years old) research subjects were imaged in an adaptive optics fluorescence lifetime imaging ophthalmoscope (AOFLIO) which has been previously described(*9*). AOFLIO images at a 1.4 degree field of view with the ability to resolve individual RPE cells were acquired. RPE fluorescence was acquired with a 532 nm excitation light and 575-725 nm collection which primarily targets lipofuscin. The fluorescence lifetime is sensitive to the environment and composition of the fluorophores. The weighted mean fluorescence lifetime (Taum) in picoseconds was computed by summing the histograms across each location and using a 2-component fit. Potentially interesting locations were chosen in the CLN3 subjects using clinical images. Reflectance images of the photoreceptors were obtained using 796 nm light.

**Recombinant Acid Ceramidase (rhAC) treatments**

Recombinant acid ceramidase (rhAC) was obtained commercially from Creative Biomart (AC-524H, Shirley, NY, USA) and/or was provided by Dr. Edward Schuchman. The rhAC contained mostly "high mannose" type oligosaccharides, and uptake studies using mouse alveolar macrophages demonstrated carbohydrate-mediated internalization of the enzyme.

The rhAC utilized in this study are internalized by mammalian cells including RPE and bioactive both in vitro (*Figure 3d and S8a ,S8b*) and *in vivo* (*Figure 8b-8d, and S14a-S14d*) (*10-12*). Furthermore, supplementation of rhAC to iRPE cells in vitro and/or intravitreal administration of rhAC in the miniswine eye impacted ceramide metabolism as measured by intracellular ceramide and S1P levels (*Figure 8b-8d, S8a, S8b and S14a-S14d*) in vitro treatments and treated with rhAC showed and has been previously utilized for rhAC administration *in vivo(11).*

**rhAC treatment of RPE monocultures.** CLN3 hESC-RPE grown on transwell membrane inserts were treated apically with 30 µg/mL dose for 1 hour at 37°C. Parallel cultures of CLN3 RPE that were not treated with rhAC were considered as untreated cultures. Following the treatments, untreated and rhAC-treated CLN3 hESC-RPE were harvested for immunocytochemistry and Western blot analyses. In a subset of experiments, untreated and rhAC-treated CLN3 hESC-RPE were fed with 40 POS/RPE cell for 30 min at 37°C with 10% FBS media supplementation. At the end of the incubation period, cells were washed with 1X PBS to remove any unbound POS from the hESC-RPE surface. Subsequently, hESC-RPE cells were harvested for Western blot analysis to measure RHO. Blots were probed for AC to confirm rhAC uptake by rhAC-treated CLN3 hESC-RPE.

**rhAC treatment of Control RO-CLN3 RPE and CLN3 RO-CLN3 RPE assembloids.** Control RO-CLN3 RPE or CLN3 RO-CLN3 RPE co-cultures on transwell inserts with TER > 150 $\Omega^*cm^{-2}$ were treated apically for a period of 7 days with rhAC (30 ug/mL dose for 1 hour), Creative Biomart, Shirley, NY, USA). Cocultures were treated daily and parallel cultures of untreated Control RO-CLN3 RPE or CLN3 RO-CLN3 RPE cocultures fed with routine culture media with daily media change served as controls in these experiments. Following the treatments, untreated and rhAC-treated assembloids were fixed for immunocytochemistry analyses.

**Intravitreal rhAC injections in CLN3 miniswine eyes.** Miniswine were pre-anesthetized with 14 mg/kg of Ketamine given intramuscularly and 1 mg/kg of acepromazine given intramuscularly, and isoflurane at 1-5% using cone inhalation delivery. Post-sedation, tropicamide 1% eyedrops was applied to both eyes for pupil dilation. The eyes were next prepped with 5% Povidone-iodine solution and a 30-32 G BD ½ or 5/16 inch needle on a tuberculin or similar syringe was be used to inject rhAC in one eye and vehicle (sterile 1X PBS) in the contralateral eye at a total maximum volume of 0.05 ml into the vitreous cavity. A sterile cotton tip applicator was held over the needle track as the needle is removed and for 5 seconds after to reduce reflux. At the completion of the procedure antibiotic/steroid drops or ointment was be applied to the eyes. Animals were euthanized day 7 post-intravitreal injection and tissue was harvested for immunohistochemical analysis. Note the rhAC used for miniswine experiment was provided by Dr. Edward Schuchman and has been previously utilized for rhAC administration *in vivo* (*11*).

**Mouse intravitreal rhAC injections and toxicity analysis**

We used 3-4 month old C57BL/6J mice (Jackson Laboratory: stock #000664) for testing the in vivo toxicity of intravitreally injected rhAC. All experiments adhered to the ARVO Statement for the Use of Animals in Ophthalmic and Vision Research and were approved by the University Committee of Animal Resources of the University of Rochester. Mice were anesthetized by intraperitoneal injection of 100mg/kg ketamine (Par Pharmaceuticals, Chestnut Ridge, NY) and 10 mg/kg xylazine (Akorn Inc, Lake Forest, IL) and pupils were dilated using an ophthalmic solution of phenylephrine 2.5% (Paragon Bioteck Inc, Portland, OR) and tropicamide 1% (Akorn Inc, Lake Forest, IL). Similar to the miniswine rhAC intravitreal administration and based on intravitreal volume of mice an equivalent dose was calculated for toxicity studies in wildtype mice. For each mouse, one eye received an intravitreal injection of 1μL of ASAH1 (0.1μg/μL) and the other eye 1μL PBS. After 1 week, retinal function was assessed using the Celeris ERG system specific for rodents (Diagnosys, Lowell MA). Prior to imagining, mice were dark-adapted for 18 hours. Dark-adapted ERG measurements were recorded at 0.01, 0.1 and 1cd.s/m$^2$ to analyze rod function, followed by light adaptation for 15 minutes and then exposure to light intensity of 3 and 10 cd.s/m$^2$ to measure cone function. Average a- and b-wave amplitudes for all eyes injected with PBS or ASAH1 were determined, and no significant differences were found between treatment groups using a Students t-test statistical analysis. Eyes were then examined by Fundus photography to look for retinal anomalies and Fluorescein angiography (FA) to visualize blood flow throughout the retina using the Micron III instrument and software (Phoenix Instruments, Naperville, IL).

**Image Analyses**

Analysis of confocal images was performed using either ImageJ or IMARIS. To determine distance between the RPE and RO in coculture images, surfaces were created using the IMARIS surfaces tool for DAPI-stained nuclei. Briefly, in the surfaces creation wizard, the DAPI channel and object-object statistics were selected, and the smooth surfaces were assigned a detail level of 5 μm with a 6.44 μm background subtraction. The threshold intensity was set such that the RPE or RO nuclei were enveloped in the surface without any of the other cell type's nuclei and the surfaces which represented the correct cell type were selected using either voxel size or z position as parameters. After surfaces representing each cell type were created, the distance between the RO and the RPE was determined by IMARIS using the parameter "shortest distance to surface X" where X is the user-assigned identity of the surface. For images where the RO and RPE layers could not be sufficiently separated by the surfaces creation wizard (distance between RO and RPE nuclei <5 μm), a distance of 5 μm was assigned. Quantification of POS in RPE was performed in ImageJ by selecting only slices which contained RPE and counting $RHO^+$ or ML-$OPSIN^+$ particles.

**Transmission Electron Microscopy (TEM) Analysis**

PEG hydrogels (Control RO-Control RPE) were fixed with 4% paraformaldehyde/2.5% glutaraldehyde in 0.1 M sodium cacodylate buffer for 24 hours prior to processing. Processing and imaging of the samples was carried out in the University of Rochester Medical Center Electron Microscope Shared Resource Laboratory.

**Transepithelial Resistance (TER) Measurements**

EVOM2 volt-ohm meter (World Precision Instruments, Sarasota, FL, USA) was used to measure TER of RPE grown in transwell inserts as described. TER measurements were reported as resistance per area or $\Omega * cm^{-2}$ after blank subtraction.

**Table S1. Abbreviations used in this paper:**

| ***Abbreviation*** | ***Meaning*** |
|---|---|
| AAV | Adeno-associated Virus |
| AC | Acid Ceramidase |
| AOFLIO | Adaptive Optics Fluorescence Lifetime Ophthalmoscopy |
| AOSLO | Adaptive Optics Scanning Light Ophthalmoscope |
| ASAH1 | N-acylsphingosine Aminohydrolase 1/Acid Ceramidase 1 |
| ASO | Antisense Oligonucleotide |
| Bp | Base Pair |
| CC | Connecting Cilium |
| COS | Cone Outer Segment |
| CRALBP | Cellular Retinaldehyde-Binding Protein |
| DMMB | 1,9-Dimethylmethylene Blue |
| DPBS | Dulbecco's Phosphate Buffered Saline |
| ECM | Extracellular Matrix |
| Ex | Exon |
| EZR | Ezrin |
| FDA | United States Food and Drug Administration |
| $^1$H NMR | Proton Nuclear Magnetic Resonance |

| | |
|---|---|
| hPSC-RPE | Retinal Pigment Epithelium Derived from Human-derived Induced Pluripotent Stem Cells |
| hPSC | Human-derived Pluripotent Stem Cell |
| hPSC-RO | Retinal Organoid Derived from Human-derived Pluripotent Stem Cells |
| hPSC-RO-RPE | 3D Retina Derived from Human-derived Pluripotent Stem Cells |
| INL | Inner Nuclear Layer |
| IPM | Interphotoreceptor Matrix |
| iPSC | Induced Pluripotent Stem Cell |
| iPSC-RO | Retinal Organoid Derived from Induced Pluripotent Stem Cells |
| JCNL | Juvenile Neuronal Ceroid Lipofuscinosis |
| kDa | Kilodalton |
| LAP | Lithium Phenyl-2,4,6-trimethylbenzoylphosphinate |
| LSD | Lysosomal Storage Disorder |
| M6P | Mannose-6-Phosphate |
| MerTK | Mer Tyrosine Kinase |
| MMP | Matrix Metalloproteinase |
| NCL | Neuronal Ceroid Lipofuscinosis |
| OCT | Optical Coherence Tomography |

| | |
|---|---|
| OLM | Outer Limiting Membrane |
| ONL | Outer Nuclear Layer |
| OS | Outer Segment |
| PEG | Poly(ethylene glycol) |
| PEG-HA | Poly(ethylene glycol) and Hyaluronan Hydrogel |
| PEG-NB | Poly(ethylene glycol)-amide-norbornene |
| PPAR-α | Peroxisome Proliferator-Activated Receptor Alpha |
| POS | Photoreceptor Outer Segment |
| RCVRN | Recoverin |
| rhAC | Recombinant Human Acid Ceramidase |
| RHO | Rhodopsin |
| RO | Retinal Organoid |
| RO-RPE | 3D Retina, Retinal Organoid-Retinal Pigment Epithelium Coculture |
| ROS | Rod Outer Segment |
| RPE | Retinal Pigment Epithelium |
| S1P | Sphingosine-1-phosphate |
| SD-OCT | Spectral Domain Optical Coherence Tomography |
| sGAG | Sulfated Glycosaminoglycan |
| TEM | Transmission Electron Microscopy |

| | |
|---|---|
| TER | Transepithelial Resistance |
| TUBB3 | Tubulin Beta 3 Class III |
| VSX2 | Visual System Homeobox 2 |
| WT | Wild-type |
| ZO-1 | Zonula Occludens-1 |

**Table S2. Primary Antibodies Used in This Paper.**

| **Primary Antibodies/stains Used in This Paper** | | | | |
|---|---|---|---|---|
| ***Antibody*** | ***Dilution for ICC*** | ***Dilution for WB*** | ***Manufacturer*** | ***Catalog Number*** |
| Mouse anti-Actin | | 1:500 | Santa Cruz | sc-47778 |
| Mouse anti-Rhodopsin | 1:100 | 1:500 | Millipore | MABN15 |
| Mouse anti-Sphingosine-1-Phosphate | 1:50 | | Echelon Biosciences | Z-P300 |
| Peanut anti-Lectin PNA-AF647 Conjugate | 1:50 | | Invitrogen | L32460 |
| Rabbit anti-ASAH1 | 1:100 | | Millipore | ABN468 |
| Rabbit anti-ASAH1 | | 1:500 | Proteintech | 11274-1-AP |
| Rabbit anti-Caveolin 1 | | 1:750 | Cell Signaling Technologies | 3267S |
| Rabbit anti-Ceramide | 1:10 | | Enzo Life Sciences | ALX-804-196-T050 |

| | | | | |
|---|---|---|---|---|
| Rabbit anti-Cleaved Caspase3-AF647 Conjugate | 1:100 | | Cell Signaling Technologies | 9602S |
| Rabbit anti-Ezrin | | 1:1000 | Cell Signaling Technologies | 3145S |
| Rabbit anti-M/L-Opsin | 1:100 | | Millipore | AB5405 |
| Rabbit anti-Phosphoezrin | | 1:500 | Invitrogen | PA5-37763 |
| Rabbit anti-Recoverin | 1:1000 | | Proteintech | 10073-1-AP |
| Sheep anti-VSX2 | 1:200 | | Exalpha | X1179P |
| Rabbit anti-CRALBP | 1:200 | | Abcam | ab15051 |
| Rabbit anti-MERTK | 1:100 | | Abcam | ab52968 |
| Rabbit anti-RPE65 | 1:100 | | GeneTex | GTX103472 |
| Mouse anti-Tubulin β 3 | 1:200 | | Biolegend | 8010201 |

| | | | | |
|---|---|---|---|---|
| Phalloidin conjugated-Alexa Fluor™ 488 | 1:100 | | ThermoFisher Scientific | A12379 |

**Supplementary material References**

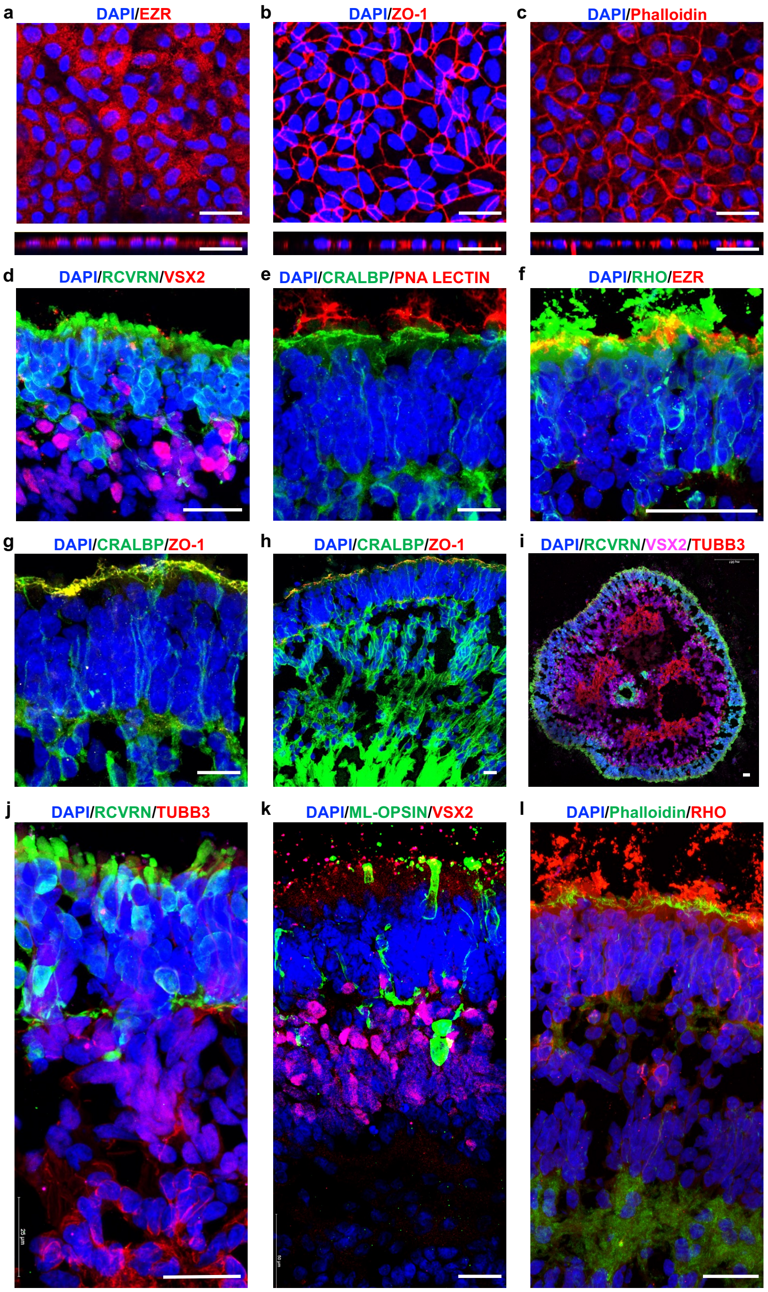


**Figure S1. Characterization of mature RPE monolayer and stage 3 retina organoid (RO) prior to use in RO-RPE assembloid culture.**

**a-c)** Immunofluorescence images of RPE wholemounts in the planar view (top panels) and orthogonal xz views (bottom panels) showing the expected localization of RPE microvilli protein, EZR, tight junction marker, ZO-1, and phalloidin-stained actin cytoskeleton. Cell nuclei are labeled with DAPI. Scale bar = 25 µm.

**d-l)** Immunofluorescence images showing the known and expected localization of RCVRN$^+$ POS (ROS and COS), RHO$^+$ ROS, ML-OPSIN and PNA-LECTIN$^+$ COS, VSX2$^+$ bipolar cells, CRALBP$^+$ Muller Glia, EZR$^+$, phalloidin$^+$, ZO1$^+$ outer limiting membrane and TUBB3$^+$ retinal ganglion cells in stage 3 RO . Scale bar = 50 µm for panels a-i and scale bar =25 µm for panels j-l.

*Related to Figure 1.*

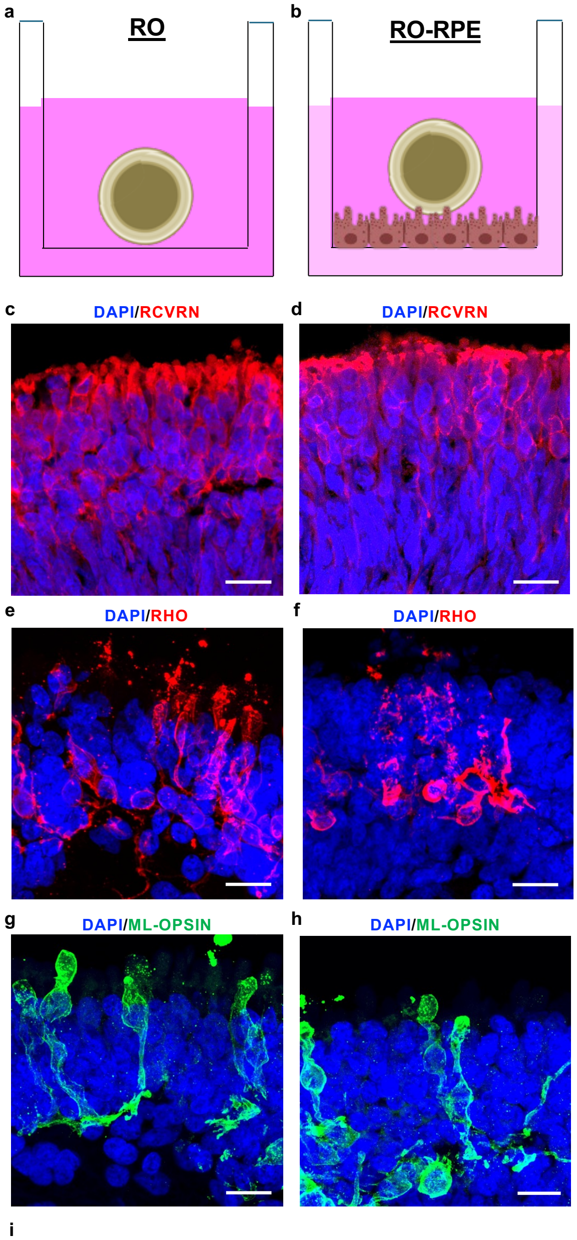


**Figure S2. Characterization of POSs at day 7 in the RO monoculture versus the RO-RPE direct co-cultures.**
**a, b)** Schematic representation of RO monocultures and RO-RPE direct co-cultures in transwells.
**c-h)** Immunofluorescence images of RO cryosections showing similar localization of RCVRN$^{+}$ POS (ROS and COS), RHO$^{+}$ ROS, ML-OPSIN$^{+}$ COS in the RO layer of the RO monocultures versus the RO-RPE co-cultures Scale bar = 15µm.
**i***Related to Figure 1.*

**Figure S3**

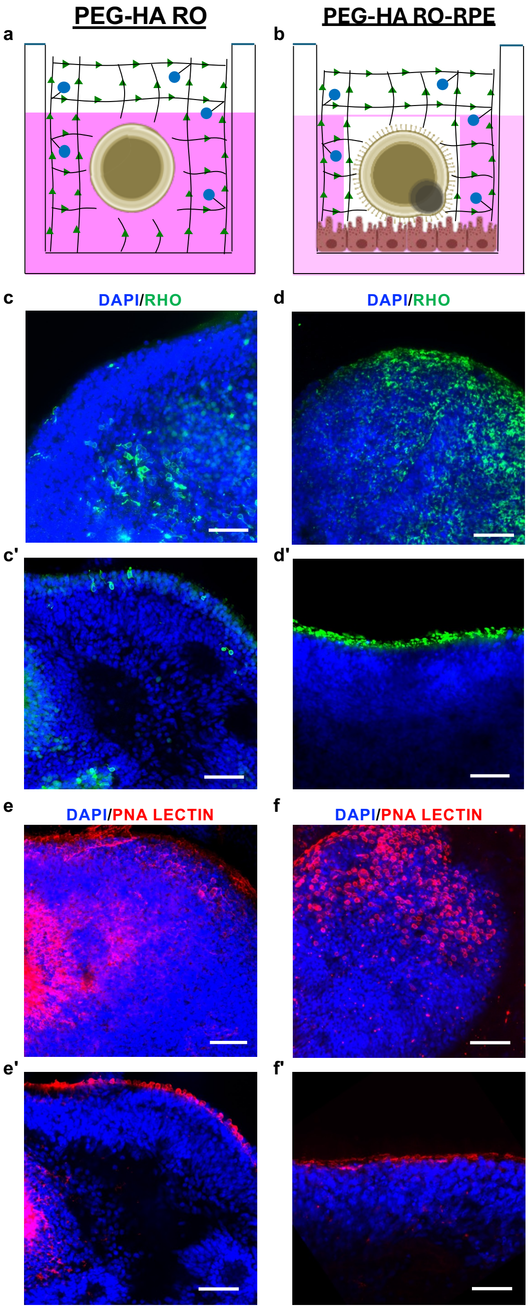


**Figure S3. Characterization of POSs in the day 7 PEG-HA RO monoculture versus the day 7 PEG-HA RO-RPE cultures .**
**a, b)** Schematic representation of the PEG-HA RO monocultures and PEG-HA RO-RPE direct co-cultures in transwells.
**c-f)** Immunofluorescence images of tissue wholemount in 3D volume view (c, d and e, f) and a single plane (c', d' and e', f') showing the improved localization of $RHO^{+}$ ROS and PNA-$LECTIN^{+}$ COS in the RO layer of PEG-HA RO-RPE cultures versus PEG-HA RO monocultures. Note that images for the RO-RPE cultures were taken on the the side of RO facing the RPE monolayer. Scale bar = 15µm. *Related to Figure 1.*

**Figure S4**

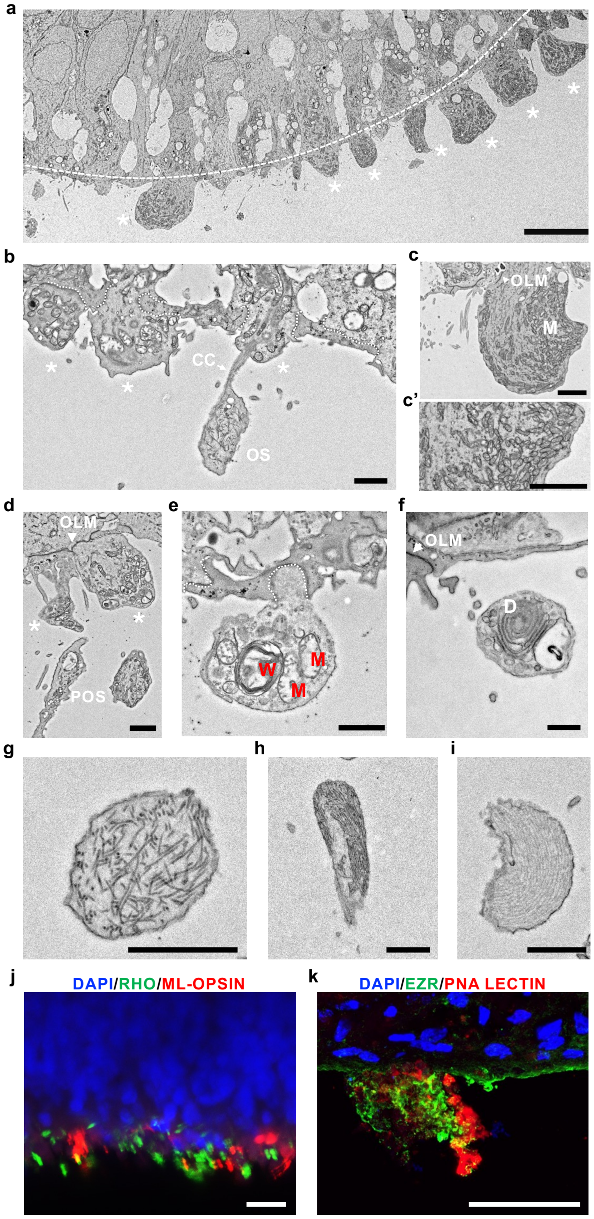


**Figure S4. Characterization of RO tissue cross-section in day 7 PEG-HA RO-RPE assembloid cultures.**

**a**) Transmission electron microscopy (TEM) image of the **of** RO tissue cross-section showing outer limiting membrane/ OLM (dashed white line) and photoreceptor inner segments (*) in the PEG-HA RO-RPE. Scale bar = 10 μm.

**b-c)** TEM image of a RO tissue cross section from the PEG-HA RO-RPE displaying photoreceptor inner segments with characteristic electron dense junctions of the OLM (dashed white line) and the coalescence of mitochondria and membranous materials (b, c, c'). The connecting cilium (CC), a hallmark feature of POS development can be seen connecting the inner segment (*) to the forming POS) featuring loosely organized membranous material bound for organization into discs (b). Higher magnification TEM image of a single photoreceptor inner segment showing the electron density of the cell-to-cell junctions of the OLM (c, c') and concentration of mitochondria (c'). Scale bar = 1 μm (panel b) and 2 μm (panel c).

**d)** TEM image of a RO tissue cross section from the PEG-HA RO-RPE showing the ultrastructure of OLM at the base of photoreceptor inner segments (*) and adjacent POS with little to no organization of discs. Note that connecting cilia are not visible in many cases this is likely a limitation of imaging 2-dimensional thin sections, 70 nm in thickness, of a 3-dimensional structure. Scale bar = 1 μm.

**e)** TEM image of a RO tissue cross section from the PEG-HA RO-RPE showing the varied composition of inner segments (*) varied, including degrading mitochondria (red M) and whirled membranes (red W) with the OLM junctions of the segment indicated by a dashed white line. Scale bar = 1 μm.

**f)** TEM image of a RO tissue cross section from the PEG-HA RO-RPE showing a POS with structures consistent with the beginning of the organization of discs that is located outside of the OLM. Scale bar = 1 μm.

**g-i)** TEM images of POSs in the RO of the PEG-HA RO-RPE showing POSs at varied maturation stages in the RO that included POSs with the disorganized formation of discs (g), POSs with the beginning of disc organization (h) and POSs with well-ordered disc stacks (i). Scale bar = 1 μm.

**j, k)** Immunofluorescence images of RO wholemount in a single plane (j) and RO-cryosection (k) showing the localization of $RHO^{+}$ ROS, ML-OPSIN+ and $PNA\text{-}LECTIN^{+}$ COS and $EZR^{+}$ OLM in the RO layer of PEG-HA RO-RPE co-cultures. Note that images for the RO-RPE cultures were taken on the the side of RO facing the RPE monolayer. Scale bar = 15μm.

*Related to Figure 1.*

**Figure S5**

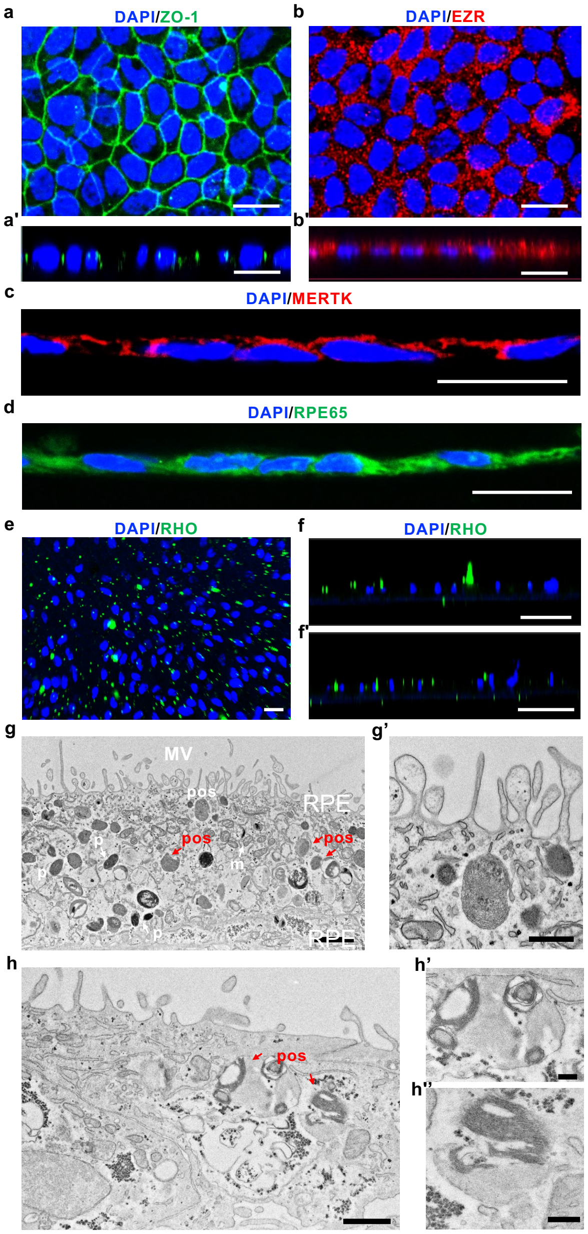


**Figure S5. Characterization of RPE wholemount and cross-section from day 7 PEG-HA RO-RPE assembloid model.**

**a,, b**) Immunofluorescence images of RPE wholemounts from the PEG-HA RO-RPE culture in the planar (a, b) and orthogonal xz views (a', b') showing the expected localization of tight junction protein, ZO-1 (green, a) and RPE microvilli protein, EZR (red, b). Cell nuclei are labeled with DAPI. Scale bar = 50 µm.

**c, d)** Immunofluorescence images of RPE cryosection from day 7 RO-RPE assembloid culture showing the expected localization of POS engulfment receptor, MERTK (red, i) and RPE signature protein, RPE65 (green, j). Scale bar = 15 µm.

**e, f)** Immunofluorescence images showing RHO+ ROS and ML-OPSIN+ COS (not included in the figure) in the RPE monolayer of day 7 RO-RPE assembloid culture in planar view (e) and orthogonal xz views (f, f') in two different planes corresponding to top (j') versus bottom (j'') of the RPE cell. DAPI (blue) indicates the presence of nuclei. Scale bar = 10 µm.

**g, g')** TEM image of a RPE tissue cross-section from day 7 PEG-HA RO-RPE cultures showing the typical apical microvilli (MV), pigmented melanosomes (p), and mitochondria (m). In addition, internalized POS at the top of the RPE monolayer (POS labeled in white, g and g') and at varied states of degradation can be seen within the RPE monolayer (POS in red). Scale bar = 1 µm (panel g) and 0.6 µm (panel Ag').

**h)** TEM image of a RPE monolayer cross-section from day 7 PEG-HA RO-RPE showing characteristic engulfed POS in the process of degradation with distinctive organized disc stack remnants. Higher magnification TEM (h' , h'') showing the ultrastructure of the degrading POS identified in (b) Scale bar = 0.8 µm (panel h) and 0.2 µm (panel h' and h'')..

*Related to Figure 1.*

Figure S6

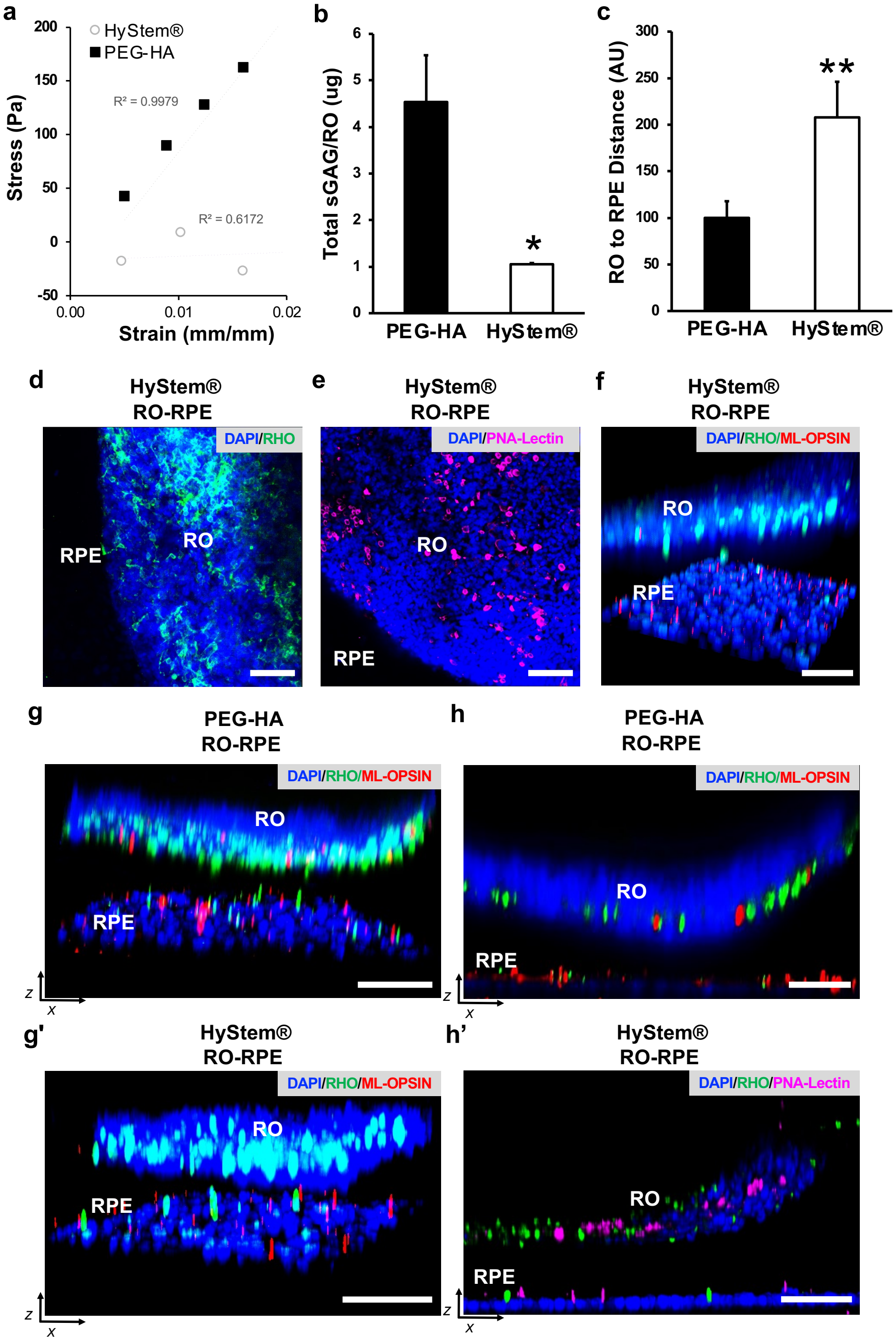


**Figure S6. Comparison of hyaluronic acid-based hydrogels in promoting tissue properties in the RO-RPE assembloid model.**

**a)** Graphical representation of elasticity properties of HyStem® and PEG-HA showing linear stress and strain relationship of PEG-HA gels but inelastic nature of HyStem® gels.

**b)** Quantitative analysis of sulfated glycosaminoglycan (sGAG) in the RO of the RO-RPE co-culture at day 7 of culture showing lower sGAG deposition in the HyStem® RO-RPE compared to the PEG-HA RO-RPE. * $p<0.05$.

**c)** Quantitative analysis of the distance between the RO and the RPE monolayer showing that RO is closer to the RPE monolayer in the PEG-HA-based RO-RPE assembloid compared to HyStem® -based RO-RPE assembloid. ** $p<0.01$.

**d-f)** Immunofluorescence images showing $RHO^+$ ROS, PNA-$Lectin^+$ COS and $DAPI^+$ cell nuclei in the RO and RPE monolayer of day 7 HyStem® RO-RPE assembloid culture in planar (d, e) and 3D orthogonal xz view (f). Scale bar = 50 µm.

**g, h)** Immunofluorescence images showing $RHO^+$ ROS, ML-$OPSIN^+$ and PNA-$Lectin^+$ COS and $DAPI^+$ cell nuclei in the RO and RPE monolayer of day 7 HyStem® versus day 7 PEG-HA RO-RPE assembloid culture in orthogonal 3D xz view (g, g') and orthogonal single-plane xz view (h, h'). Scale bar = 50 µm.

Biological replicate n ≧ 3 for all Figure S1 experiments.

*Related to Figure 1.*

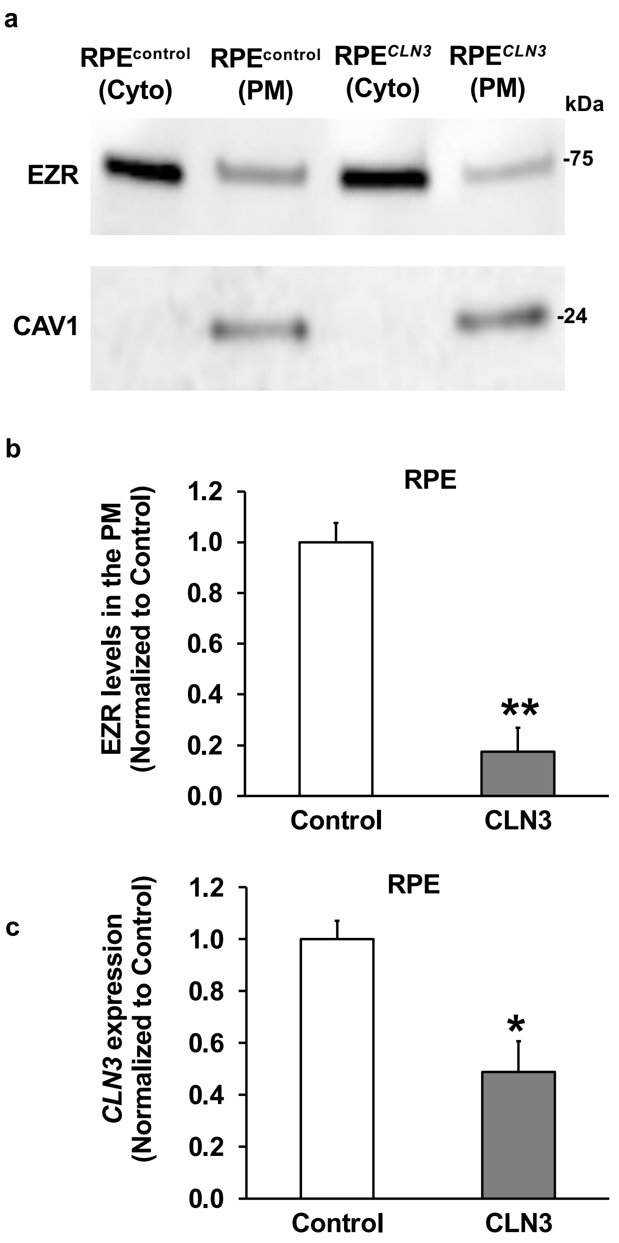


**Figure S7. Molecular changes in RPE$^{\text{control}}$ versus RPE$^{\mathit{CLN3}}$ monocultures.**

**a, b)** Western blot images (a) and quantitative analyses (b) showing altered levels of RPE microvilli protein, EZR, and plasma membrane protein, CAV1, in the cytosolic (Cyto) and plasma membrane (PM) fraction of RPE$^{\text{control}}$ versus RPE$^{\mathit{CLN3}}$ monocultures post-subcellular fractionation. Data is presented normalized to control samples. ** $p<0.01$.

**c)** Quantitative analysis of real-time RT-PCR data showing lower *CLN3* gene expression in monocultures of RPE$^{\mathit{CLN3}}$ *compared to monoculture of* RPE$^{\text{control}}$ cells. *18S* was used as the housekeeping gene and data are presented relative to *18S* and normalized to control sample. * $p<0.05$.

*Related to Figure 2-8.*

Figure S8

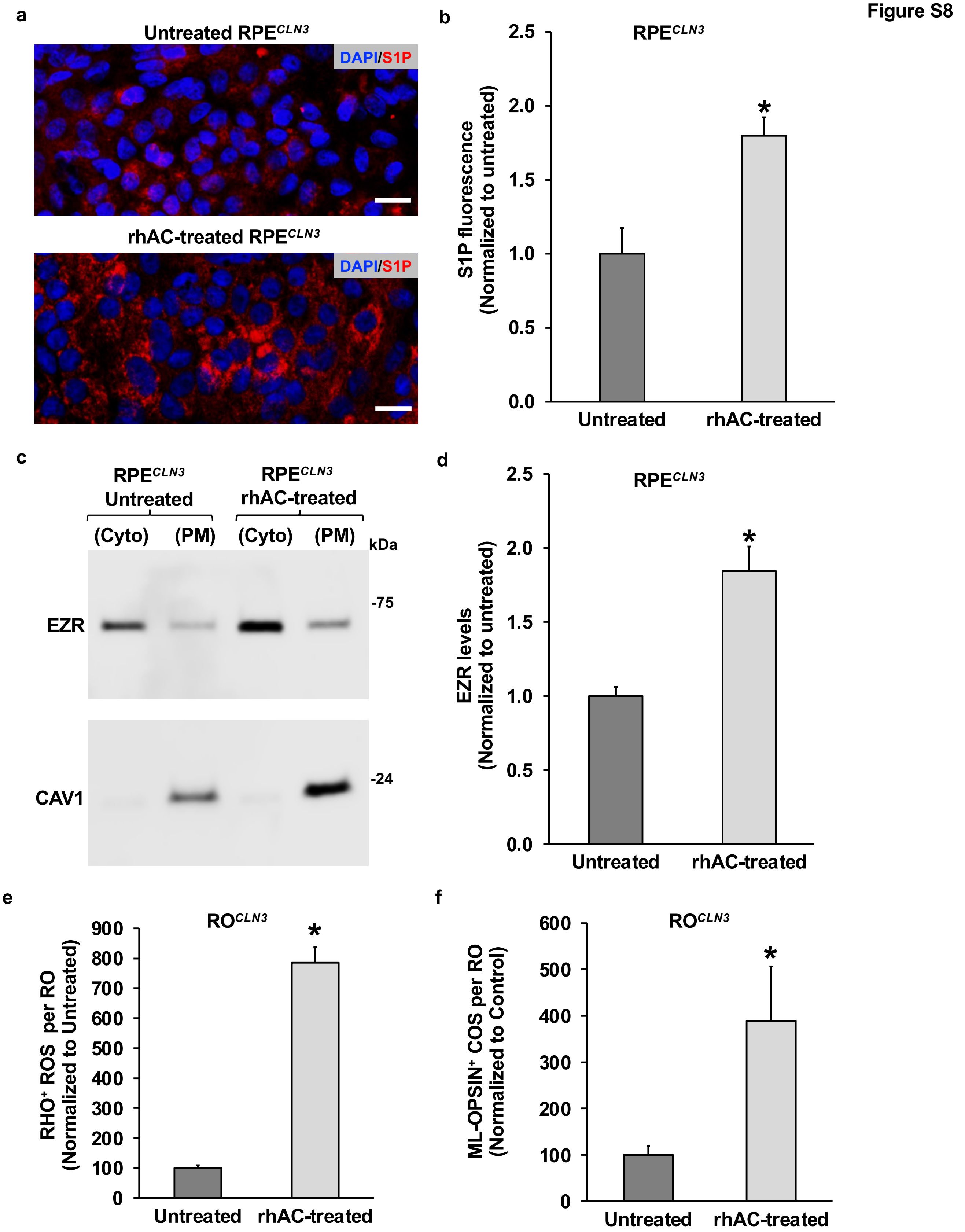


**Figure S8. impact of rhAC supplementation on S1P and EZR levels in *RPE$^{CLN3}$* monocultures.**

**a, b)** Immunofluorescence images (a) and quantitative analysis (b) illustrating the increased levels of S1P fluorescence (red, a) in rhAC-treated RPE$^{CLN3}$ compared to untreated RPE$^{CLN3}$ monocultures. S1P fluorescence intensity was measured per viewing area and data is presented normalized to untreated RPE culture (b). DAPI stained cell nuclei are shown in blue (a). Scale bar = 50 µm. * $p<0.05$.

**c, d)** Western blot images (c) and quantitative analyses (d) showing levels of EZR and CAV1, in the Cyto and PM fraction of untreated RPE$^{CLN3}$ versus rhAC-treated RPE$^{CLN3}$ monocultures post-subcellular fractionation.

**e, f)** Quantitative analyses of RHO$^{+}$ ROS (e) and ML-OPSIN$^{+}$ COS (f) in the RO layers of the untreated RO$^{control}$-RPE$^{CLN3}$ versus rhAC-treated RO$^{control}$-RPE$^{CLN3}$ cultures. Data is presented normalized to control samples. * $p<0.05$, ** $p<0.01$.

*Related to Figure 2-8.*

Figure S9

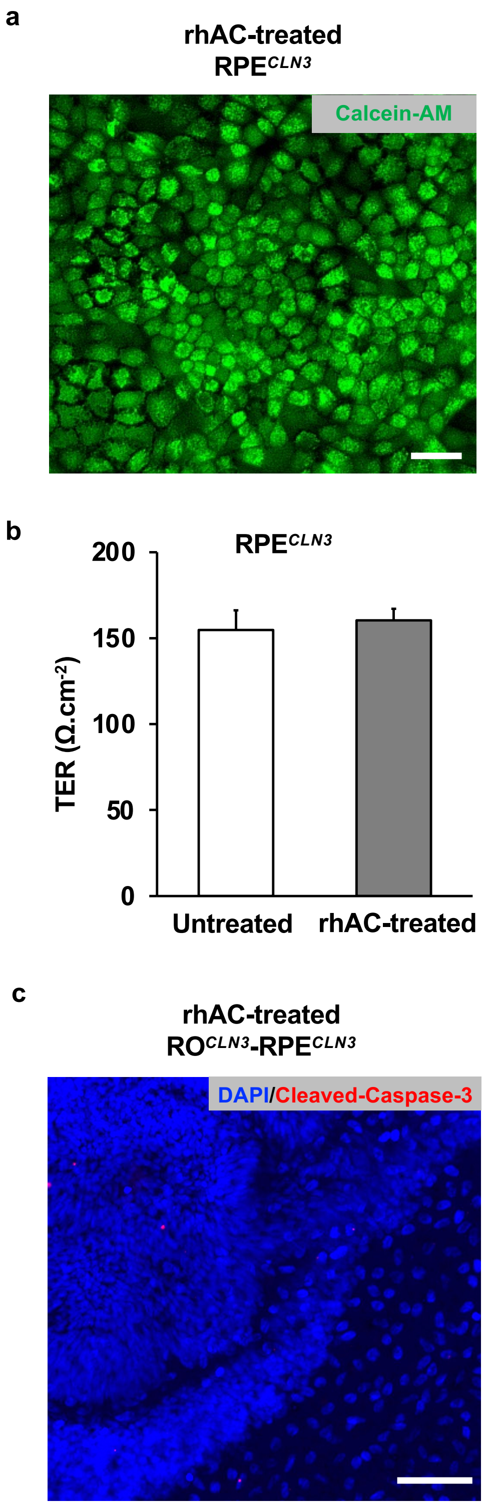


**Figure S9. No adverse impact of rhAC supplementation on on RPE$^{CLN3}$ monocultures and RO$^{CLN3}$-RPE$^{CLN3}$ assembloid cultures.**
**a-c)** Light microscopy image (a), transepithelial resistance (TER) recording (b) and immunofluorescence image (c) showing Calcein-AM$^{+}$ nuclei in live cells (green, a), TER measurements (b) and cleaved-caspase 3$^{+}$ cells (red, c) in RPE$^{CLN3}$ monocultures (a, b) and/or RO$^{CLN3}$-RPE$^{CLN3}$ co-cultures (c) after 7 days of daily rhAC treatment. TER measurement (b) is shown in comparison to untreated RPE$^{CLN3}$ monocultures. Cell nuclei were labeled with DAPI (blue, c). Scale bar = 50 µm.
Biological replicate n ≧ 3 for all Figure S2 experiments.
*Related to Figure 2-8.*

Figure S10

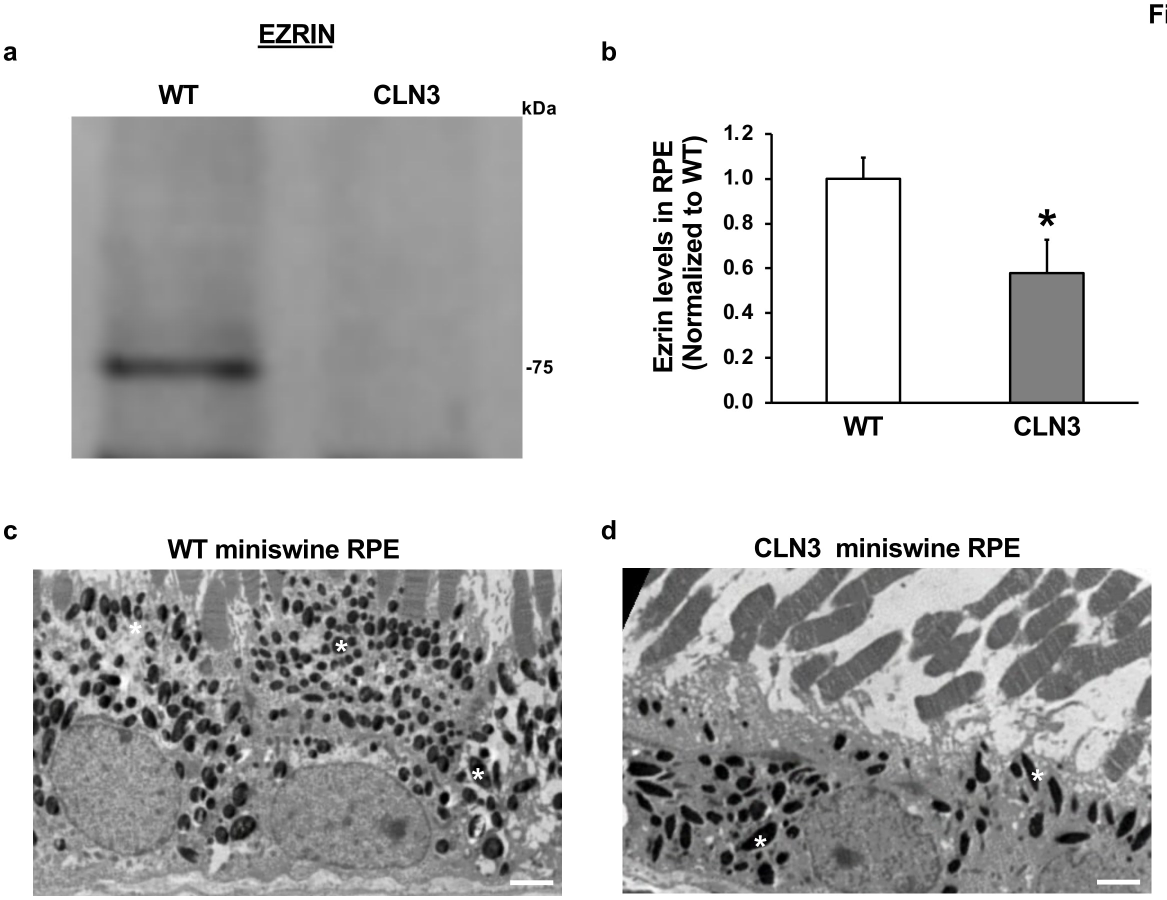


**Figure S10. Reduced levels of EZR and pigment granules in *CLN3* miniswine RPE .**

**a, b)** Western blot image (a) and quantitative analyses (b) showing reduced levels of EZR in wild-type (WT) versus CLN3 miniswine RPE at 1-month of age.

**c, d)** Electron microscopy images showing the expected apical localization of melanosomes (pigment granules, *) in wild-type (WT) miniswine RPE. In contrast, melanosomes in the *CLN3* miniswine RPE are unevenly distributed in the cytoplasm and accumulate along the basal surface Scale bar = 2 µm.

Biological replicate n ≧ 3 for all Figure S2 experiments.

*Related to Figure 4-6 and 8.*

Figure S11

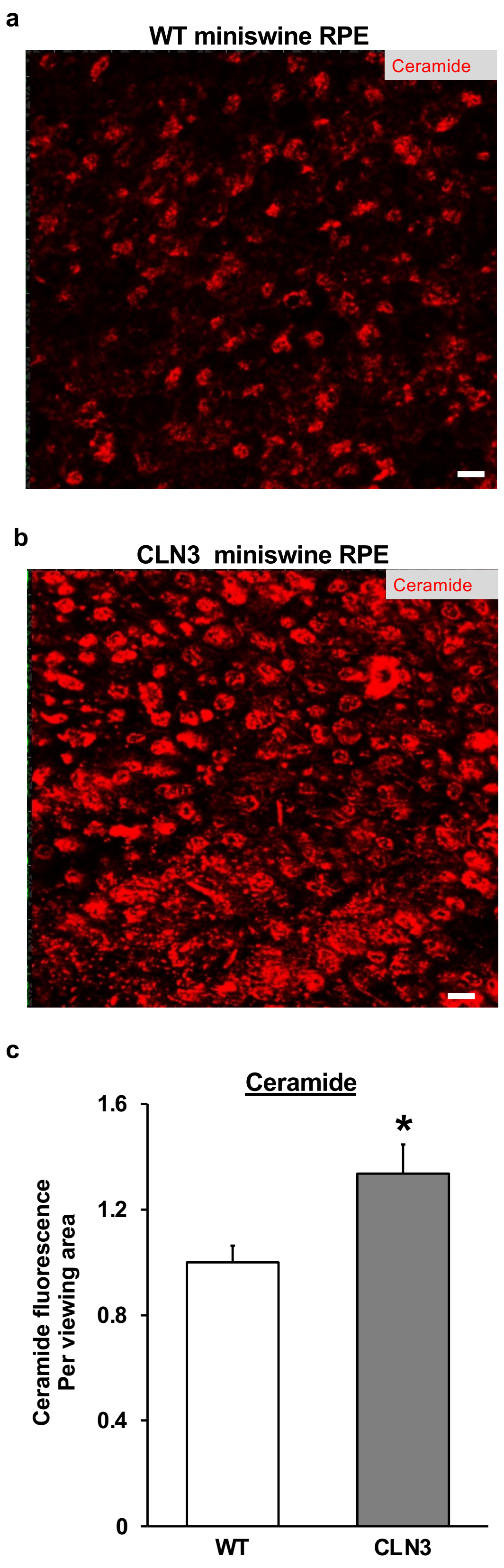


**Figure S11. Comparison of total ceramide levels in wild-type and CLN3 miniswine retina.**

**a, b)** Immunofluorescence images showing total ceramide staining in the wild-type (WT) and CLN3 miniswine retina. Cell nuclei are labeled with DAPI (blue). Scale bar = 50 µm.

**c)** Quantitative analyses of ceramide fluorescence intensity (per viewing area) in the wild-type (WT) and CLN3 miniswine retina wholemounts. * $p<0.05$.

Biological replicate n ≧ 3 for all Figure S3 experiments.

*Related to Figure 4-6 and 8.*

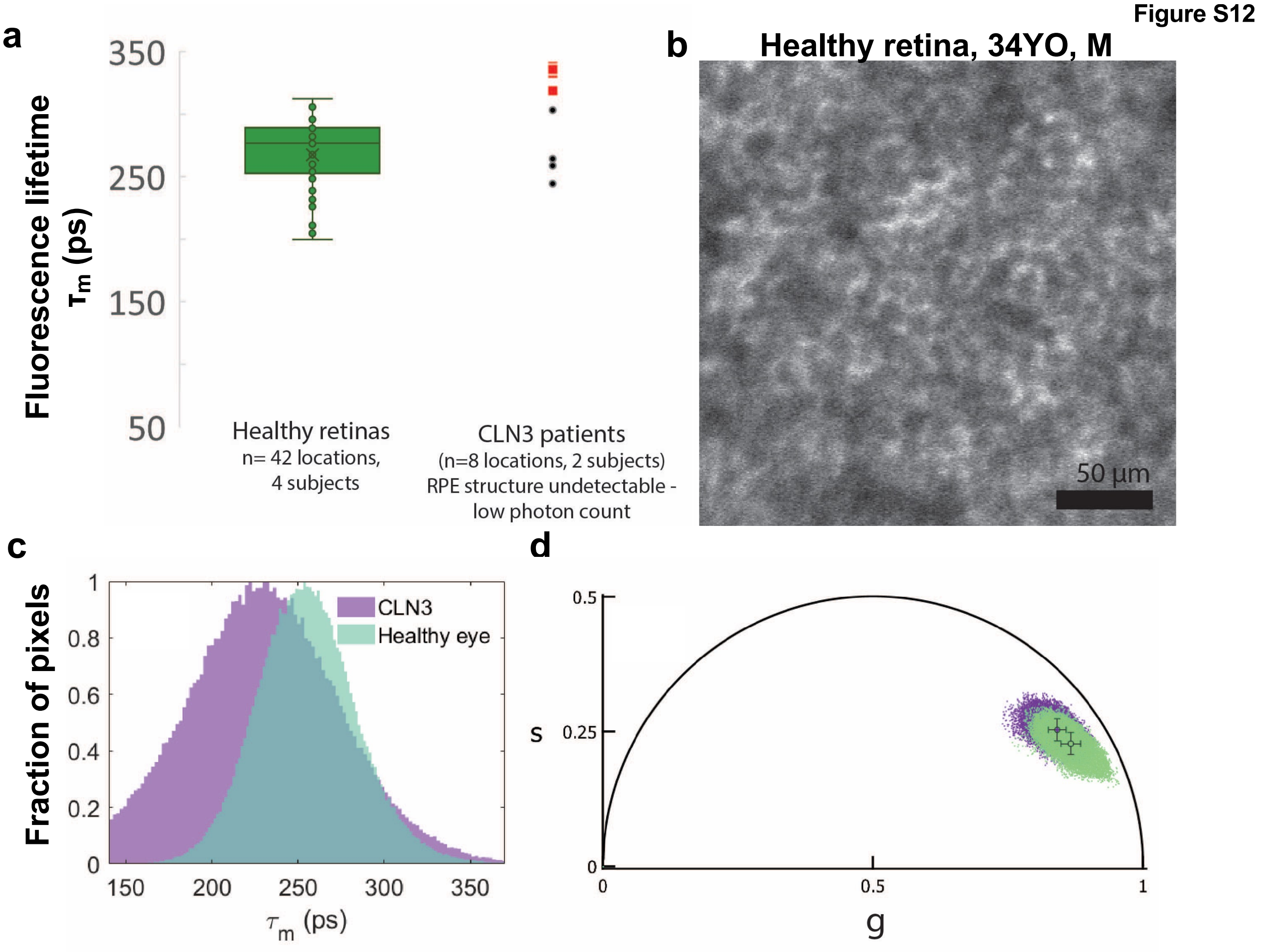


**Figure S7. Measurement of RPE lipofuscin with adaptive optics fluorescence lifetime ophthalmoscopy (AOFLIO).**

**a)** AOFLIO measurement of mean fluorescence lifetime in healthy retinas and the CLN3 disease patient retina. The fluorescence lifetime measurement indicated that the composition of measured fluorophores in healthy retina and CLN3 disease retina is very similar; fluorescence lifetime measurements at 532 nm is primarily influenced by lipofuscin (J. A. H. Tang et al., "Characterizing and identifying the fluorescence lifetime of the in vivo human RPE cellular mosaic," University of Rochester (2023).). This normative data has been previously published with a different analysis method in (J. A. H. Tang et al., "Characterizing and identifying the fluorescence lifetime of the in vivo human RPE cellular mosaic," University of Rochester (2023). and J. A. H. Tang et al., "Adaptive optics fluorescence lifetime imaging ophthalmoscopy of in vivo human retinal pigment epithelium," Biomed. Opt. Express 13(3), 1737–1754 (2022).) The CLN3 data is novel to this paper.

**b)** AOFLIO measurement of fluorescence intensity revealing the RPE structure in a healthy retina. The RPE structure in the CLN3 patient retina was undetectable most likely due to the low photon count due to a decrease in RPE autofluorescence (lipofuscin).

**c, d)** Histograms of AOFLIO measurements of the fluorescence lifetimes in a CLN3 disease retina and an age-matched healthy retina (c). The phasor plot shows similar spread of fluorophore composition in the fluorescence lifetime data between CLN3 disease retina and an age-matched healthy retina.

N=2 CLN3 disease patients.

*Related to Figure 6*

Figure S13

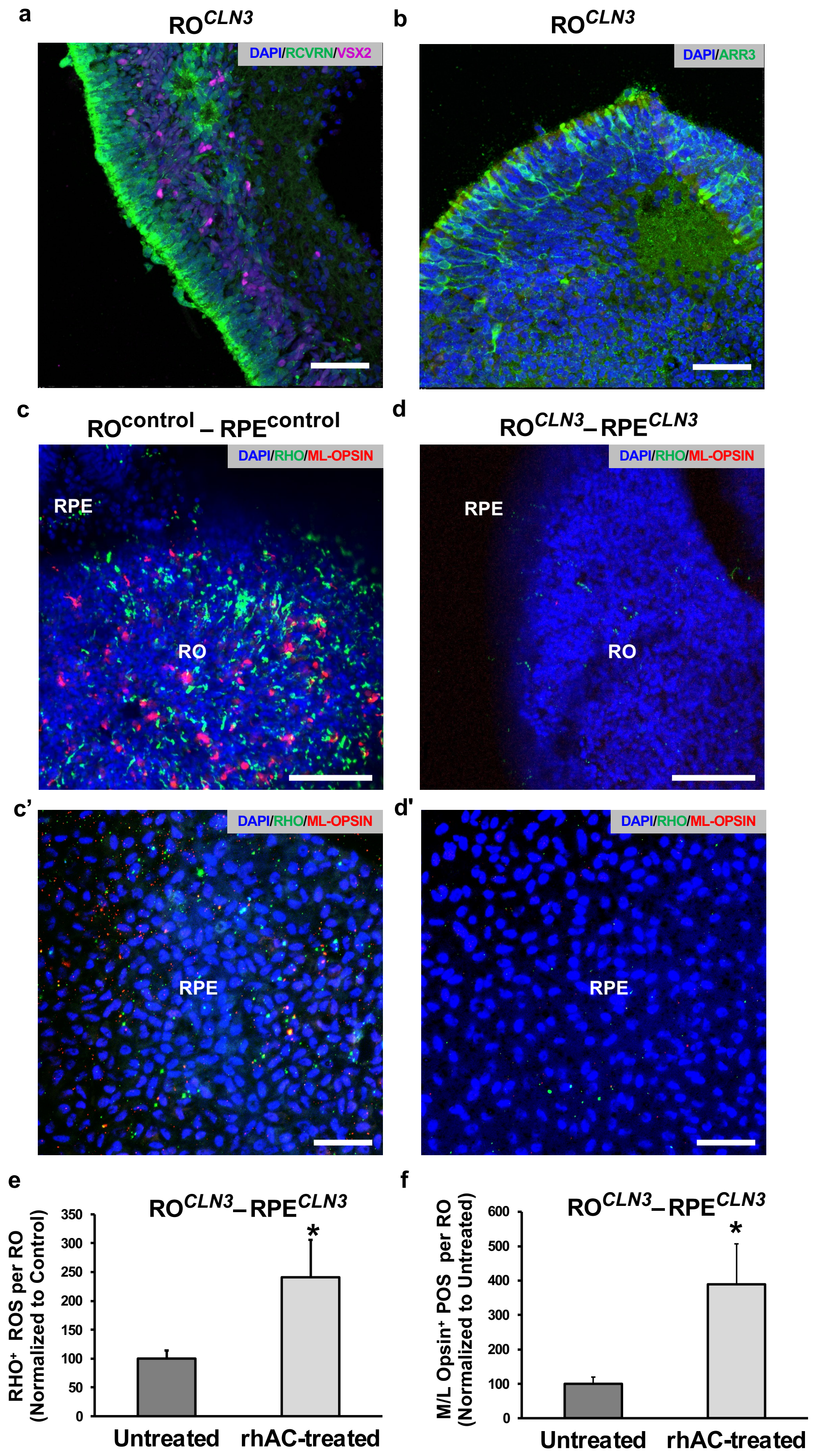


**Figure S13. RO$^{CLN3}$ -RPE$^{CLN3}$ model shows reduced levels of POS in the RO and RPE layer.**

**a)** Immunofluorescence images of RO$^{CLN3}$ showing presence of RCVRN$^{+}$ photoreceptors (a) VSX2$^{+}$ bipolar cells and DAPI$^{+}$ cell nuclei in RO$^{CLN3}$ monocultures. Scale bar = 50 µm.

**b)** Immunofluorescence images illustrating Arrestin 3 (ARR3)$^{+}$ COS and DAPI$^{+}$ cell nuclei in RO$^{CLN3}$ monocultures. Scale bar = 50 µm.

**c, d)** Immunofluorescence images of RHO$^{+}$ ROS and M/L Opsin$^{+}$ COS in the RO-RPE (c, d) and RPE monolayer (c', d') of RO$^{control}$-RPE$^{control}$ (c, c') and RO$^{CLN3}$ -RPE$^{CLN3}$ (d, d') assembloids at day 7 of culture. Images are shown in planar view and cell nuclei are labeled with DAPI.

**e, f)** Quantitative analyses of RHO$^{+}$ ROS (e) and ML-OPSIN$^{+}$ COS (f) in the RO layer of the untreated RO$^{CLN3}$ -RPE$^{CLN3}$ RPE vs rhAC-treated RO$^{CLN3}$ -RPE$^{CLN3}$ RPE assembloid cultures. Scale bar = 50 µm. * $p<0.05$.

Biological replicate n ≧ 3 for all Figure S13 experiments.

*Related to Figure 7.*

**Figure S14**

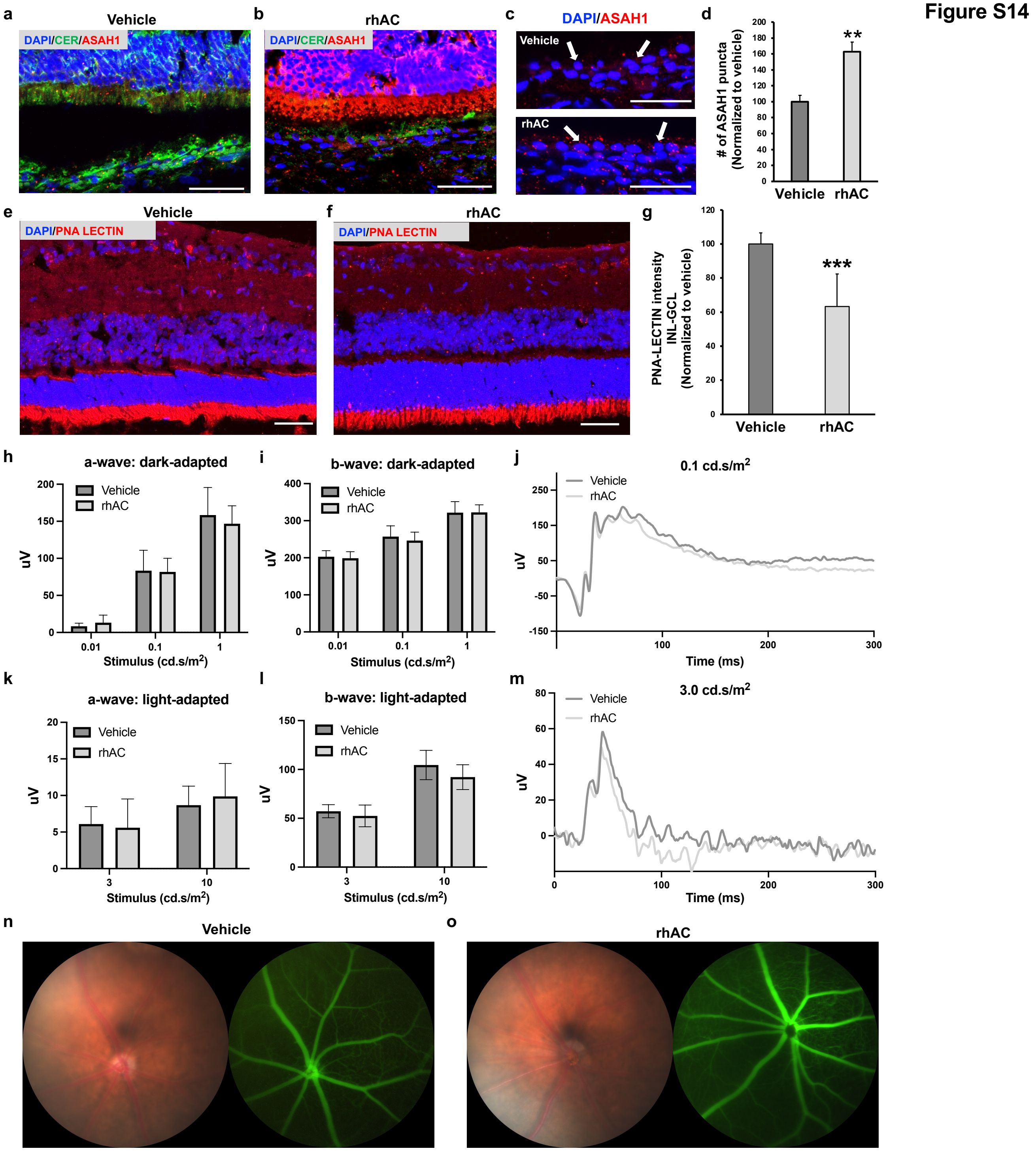


**Figure S14. Characterization of rhAC efficacy and toxicity in the retina *in vivo*.**

**a, b)** Immunofluorescence images of retina cryosections showing increased ceramide (green) and decreased ASAH1 (red) levels in vehicle injected (a) versus rhAC-injected (b) CLN3 miniswine photoreceptor, RPE and choroid at day 7 of treatment. Cell nuclei (DAPI, blue). Scale bar = 50 µm.

**c, d)** Immunofluorescence images of RPE-choroid cryosections showing expected ASAH1 localization (red, c) and quantitative analyses of showing increased ASAH1 levels in the RPE monolayer (d) in vehicle injected versus rhAC-injected (b) CLN3 miniswine eye at day 7 of treatment. Cell nuclei (DAPI, blue). RPE monolayer is demarcated by a white arrow. Scale bar = 50 µm.

**e-g)** Immunofluorescence images of retina cryosection (e, f) and quantitative analyses showing improved PNA-LECTIN localization (e, f) with reduced intensity of PNA-LECTIN in the inner retina inner nuclear layer/ INL- ganglion cell layer/GCL) (g) in rhAC-injected CLN3 miniswine eye compared to vehicle injected versus at day 7 of treatment. Cell nuclei (DAPI, blue). Scale bar = 50 µm.

**h-m)** Graphical illustration (h, i,, k and l) and representative ERG traces (j, m) showing similar a-wave and b-wave electroretinogram (ERG) amplitudes in dark-adapted (h-j) and light-adapted conditions (k-m) of vehicle-injected versus rhAC-injected C57BL/6J mouse eyes at day 7 post-injection.

**n, o)** Fundus photograph and fluorescein angiography images of vehicle-injected versus rhAC-injected C57BL/6J mouse eyes at day 7 post-injection showing no adverse effect of intravitreally injected rhAC on the mouse retina.

Biological replicate $n \geqq 3$ for all Figure S14 experiments

*Related to Figure 8*

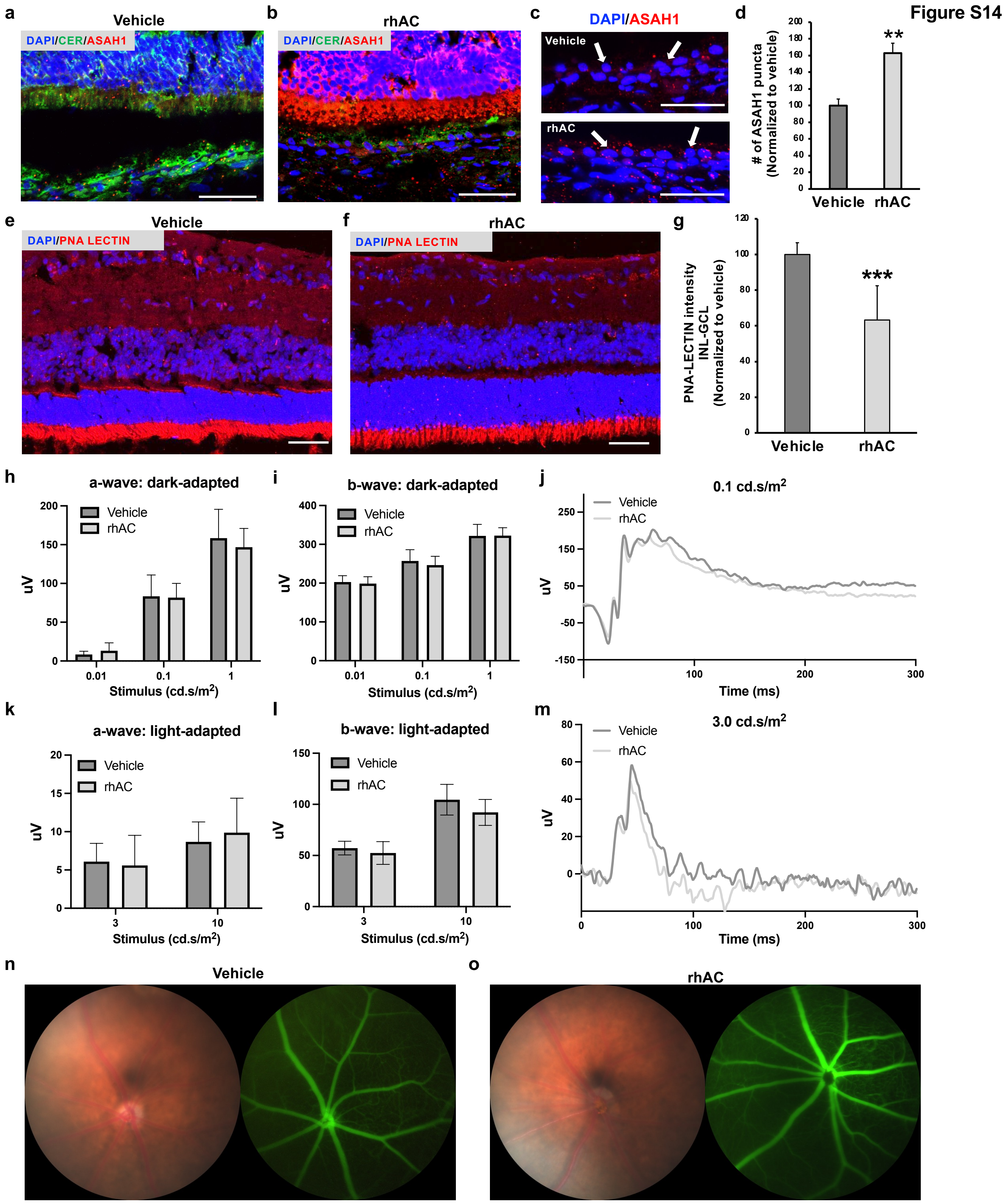


**Figure S14. Characterization of rhAC efficacy and toxicity in the retina *in vivo*.**

**a, b)** Immunofluorescence images of retina cryosections showing increased ceramide (green) and decreased ASAH1 (red) levels in vehicle injected (a) versus rhAC-injected (b) CLN3 miniswine photoreceptor, RPE and choroid at day 7 of treatment. Cell nuclei (DAPI, blue). Scale bar = 50 µm.

**c, d)** Immunofluorescence images of RPE-choroid cryosections showing expected ASAH1 localization (red, c) and quantitative analyses of showing increased ASAH1 levels in the RPE monolayer (d) in vehicle injected versus rhAC-injected (b) CLN3 miniswine eye at day 7 of treatment. Cell nuclei (DAPI, blue). RPE monolayer is demarcated by a white arrow. Scale bar = 50 µm.

**e-g)** Immunofluorescence images of retina cryosection (e, f) and quantitative analyses showing improved PNA-LECTIN localization (e, f) with reduced intensity of PNA-LECTIN in the inner retina inner nuclear layer/ INL- ganglion cell layer/GCL) (g) in rhAC-injected CLN3
